# Regulation of the Drosophila transcriptome by Pumilio and CCR4-NOT deadenylase

**DOI:** 10.1101/2023.08.29.555372

**Authors:** Rebecca J. Haugen, Catherine Barnier, Nathan D. Elrod, Hua Luo, Madeline K. Jensen, Ping Ji, Craig A. Smibert, Howard D. Lipshitz, Eric J. Wagner, P. Lydia Freddolino, Aaron C. Goldstrohm

## Abstract

The sequence-specific RNA-binding protein Pumilio controls development of *Drosophila*; however, the network of mRNAs that it regulates remains incompletely characterized. In this study, we utilize knockdown and knockout approaches coupled with RNA-Seq to measure the impact of Pumilio on the transcriptome of *Drosophila* cells. We also used an improved RNA co-immunoprecipitation method to identify Pumilio bound mRNAs in *Drosophila* embryos. Integration of these datasets with the content of Pumilio binding motifs across the transcriptome revealed novel direct Pumilio target genes involved in neural, muscle, wing, and germ cell development, and cellular proliferation. These genes include components of Wnt, TGF-beta, MAPK/ERK, and Notch signaling pathways, DNA replication, and lipid metabolism. Additionally, we identified the mRNAs regulated by the CCR4-NOT deadenylase complex, a key factor in Pumilio-mediated repression, and observed concordant regulation of Pumilio:CCR4-NOT target mRNAs. Computational modeling revealed that Pumilio binding, binding site number, density, and sequence context are important determinants of regulation. Moreover, the content of optimal synonymous codons in target mRNAs exhibits a striking functional relationship to Pumilio and CCR4-NOT regulation, indicating that the inherent translation efficiency and stability of the mRNA modulates their response to these trans-acting regulatory factors. Together, the results of this work provide new insights into the Pumilio regulatory network and mechanisms, and the parameters that influence the efficacy of Pumilio-mediated regulation.

## INTRODUCTION

Eukaryotic transcriptomes are regulated by RNA-binding proteins (RBPs) that control processing, localization, stability, and - for mRNAs - translation. Pumilio proteins are one such RBP class that regulate specific mRNAs in the cytoplasm. In metazoans, Pumilio orthologs are essential for development and control of cellular proliferation and stem cell differentiation (Arvola et al. 2017; Goldstrohm 2018). Their dysfunction contributes to diseases such as the neurodegenerative disease spinocerebellar ataxia (SCA47), infertility, and cancer (Gennarino et al. 2015; Gennarino et al. 2018; Goldstrohm 2018). To better investigate the regulatory roles played by Pumilio proteins in an accessible and well-studied model system, here we focus on *Drosophila melanogaster* Pumilio (Pum), which controls germline stem cell proliferation, embryonic development, and neurological functions (Arvola et al. 2017). Pum serves as an archetype for understanding post-transcriptional regulatory mechanisms and their impact on the transcriptome.

Pumilio proteins are defined by a conserved RNA-binding domain with eight repeated triple alpha-helical units. Their RNA recognition mechanism is well understood, with each repeat presenting three amino acids that specifically contact a ribonucleotide base (Wang et al. 2002; Weidmann et al. 2016). The specificity, affinity, and 3D structure of *Drosophila* Pum bound to RNA have been extensively characterized, thereby defining the Pumilio Response Element (PRE) with consensus 5′-UGUANAUA (where N = A,G,C, or U) (Zamore et al. 1997; Zamore et al. 1999; Gerber et al. 2006; Laver et al. 2015; Weidmann et al. 2016).

Pum is a repressor that reduces translation and stability of select target mRNAs (Arvola et al. 2017). The PRE is necessary and sufficient to confer Pum-mediated regulation when placed into the 3′UTR of a reporter gene (Weidmann and Goldstrohm 2012). In this context, Pum represses protein expression by accelerating mRNA degradation and by antagonizing the translation-promoting activities of the poly(A) tail and poly(A) binding protein (Weidmann and Goldstrohm 2012; Weidmann et al. 2014; Burow et al. 2015; Arvola et al. 2020).

While Pum-regulated mRNAs have been identified by genetic and biochemical analyses (reviewed by Arvola 2017), the potential impact of *Drosophila* Pum on the transcriptome remains to be fully characterized. Robust functional evidence is limited to a handful of key Pum target genes linked to phenotypes in embryos (e.g. *hunchback*)(Murata 95, Wreden 97, Forbes 1998, Wharton 1998), the germline (e.g. *Cyclin B, mei-P26*)(Kadyrova 2007, Joly 2013), and brain (e.g. *paralytic, hid*)(Mee, 2004; Bhogal 2016). Two lines of evidence suggest a broader regulatory role of Pum. First, PRE motifs are prevalent in the transcriptome, with over 3724 genes having one or more PREs, which are distributed in 5′ UTR, 3′ UTR, and protein coding sequence (CDS)(**Table S1**)(Arvola 2017). Second, hundreds of PRE-containing mRNAs that co-immunoprecipitate with Pum from embryos or adult ovaries were identified by microarray analyses (Laver 2015, Gerber 2006). Nevertheless, regulation of these PRE-containing mRNAs by Pum remains largely unexplored.

Analysis of PRE-containing reporter genes established that Pum accelerates mRNA degradation by binding and recruiting the CCR4-NOT deadenylase complex (Weidmann et al. 2014; Arvola et al. 2020). CCR4-NOT catalyzes removal of the 3′ poly-adenosine tail of mRNAs. The poly(A) tail can stabilize mRNAs and promote their translation (Passmore and Coller 2021). Removal of the poly(A) by deadenylases initiates mRNA decay pathways (Meyer et al. 2004; Temme et al. 2004; Goldstrohm and Wickens 2008; Temme et al. 2010; Temme et al. 2014). In vivo, several Pumilio target mRNAs were shown to be degraded by CCR4-NOT (Wreden et al. 1997; Joly et al. 2013; Arvola et al. 2017; Arvola et al. 2020). However, the impact of the Pum:CCR4-NOT regulatory mechanism on the fly transcriptome has not yet been fully investigated. Moreover, the broader effects of CCR4-NOT on transcript levels in *Drosophila,* and the extent to which its activity occurs through interactions with Pum versus other factors, is not well-understood. Across eukarya, CCR4-NOT mediated deadenylation has emerged as a crucial node for mRNA regulation by multiple RNA-binding proteins and microRNAs (Goldstrohm and Wickens 2008; Jonas and Izaurralde 2015; Raisch and Valkov 2022). In addition, CCR4-NOT is implicated in codon optimality mediated mRNA decay, based on evidence that yeast CCR4-NOT interacts with ribosomes whose elongation is slowed by suboptimal codon content (Buschauer et al. 2020; Bae and Coller 2022; Wu and Bazzini 2023). The relationship between CCR4-NOT activity and codon optimality in metazoans remains to be examined. Additionally, the potential relevance of codon optimality to Pum-mediated mRNA decay is unknown.

To better understand the Pum regulatory network, both in terms of its targets and mechanisms of action, we measured the impact of Pum on the transcriptome using RNA-Seq combined with two loss of function strategies: transient depletion and knockout. Integrative analysis of the resulting differential expression data reveals a collection of PRE-enriched, Pum-bound, directly regulated target mRNAs with key functions in neural and germ cell development, transposon suppression, metabolism, cell proliferation, and signaling. Our analysis provides insights into the efficacy of Pum-mediated repression in relation to the location, sequence context, number, and density of PREs. We also measure the effect of CCR4-NOT depletion on the transcriptome. The intersection of Pum and CCR4-NOT regulated mRNAs supports the role of the deadenylase complex in regulation of natural Pum target mRNAs. Analysis of the mRNAs regulated by Pum and CCR4-NOT identified both shared and unique functional classes of target genes related to developmental pathways and potential cis-acting regulatory features. Strikingly, optimal synonymous codons exhibited a robust functional relationship to regulation by both Pum and CCR4-NOT. Collectively, our results provide new insights into the Pum regulatory network and mechanisms.

## MATERIALS AND METHODS

### Cell culture

This study utilized *Drosophila* Line 1 (DL1) cells (Drosophila Genomics Resource Center)(Schneider 1972), which were grown at 25°C in Schneider′s Drosophila medium (SDM, Gibco) supplemented with glutamine (1x GlutaMAX, Gibco), 1x antibiotic-antimycotic containing 100 units/mL of penicillin, 100 µg/mL of streptomycin, and 0.25 µg/mL of Amphotericin B (Thermo Fisher), and 10% heat inactivated fetal bovine serum (FBS, GenClone).

### Generation of DL1 Pum-myc and V5-Raf cell lines

A myc epitope tag was engineered onto the C-terminus of the *pumilio* (FBgn0003165) coding region using CRISPR-Cas9 and homologous recombination in the DL1 cell line. A guide RNA site targeting exon 13 was identified using Benchling CRISPR RNA guide software. The single guide RNA plasmid pAc sgRNA Cas9 Not1 was created by inserting annealed primers RJH284 Pum exon 13 sg1 Fwd 5′-ttcgAGGAAATAACAAATTAAGCC and RJH285 Pum exon 13 sg1 Rev 5′-aacGGCTTAATTTGTTATTTCCTc into the BspQ1 site (Bassett et al. 2013; Haugen et al. 2022). Note that the 5′ “ttc” or “aac” (in lowercase) are BsqQI cohesive overhangs and the g:c base pair was included as part of the U6 promoter. To integrate the tag, a single-stranded homology directed repair donor template (IDT) containing *Pumilio* homology arms (in capital letters), a cleavage site for the human rhinovirus (HRV) 3C protease (in lower case italics), a myc epitope tag (lower case underlined), and in-frame stop codon (bold, capital letters) was created with the following sequence: Pum-myc ssODN: 5′-ACAGCAGCTTGGGTCCCATTGGACCCCCGACCAACGGCAACGTTGTG*ctggaggtgctgttccagg gcccc*<u>gaacaaaaactcatctcagaagaggatctg</u>**TAA**AGGAAATAACAAATTAAGCCAAGCAGTCAAAGG AAACTTCTTTCTCGAATCGCAGTATAGTTTTTAGAAGCTGTAGAGCTTAACATAAACAACAAG. DL1 cells (2 x 10^6^ per well) were plated in 2 mL of SDM in one well of a 6-well plate. After 24 hours, cells were transfected with 4 µL FuGene HD (Promega), 1 µg sgRNA-Cas9 plasmid DNA, 40 pmol ssODN template, and 100 µL serum free SDM. After incubation for 48 hours at 25°C, the medium was replaced with fresh SDM containing 5 µg/mL puromycin (Gibco). 72 hours later, when cells approached confluence, they were expanded into a 10 cm dish and allowed to continue growing to 80% density. Clonal lines were then isolated by limiting dilution. The Pum-myc cell line was identified by western blot detection with anti-myc (**Figure 1**) and confirmed by PCR amplification and sequencing of Pum exon 13 from genomic DNA (**Figure S1**).

**Figure 1.**
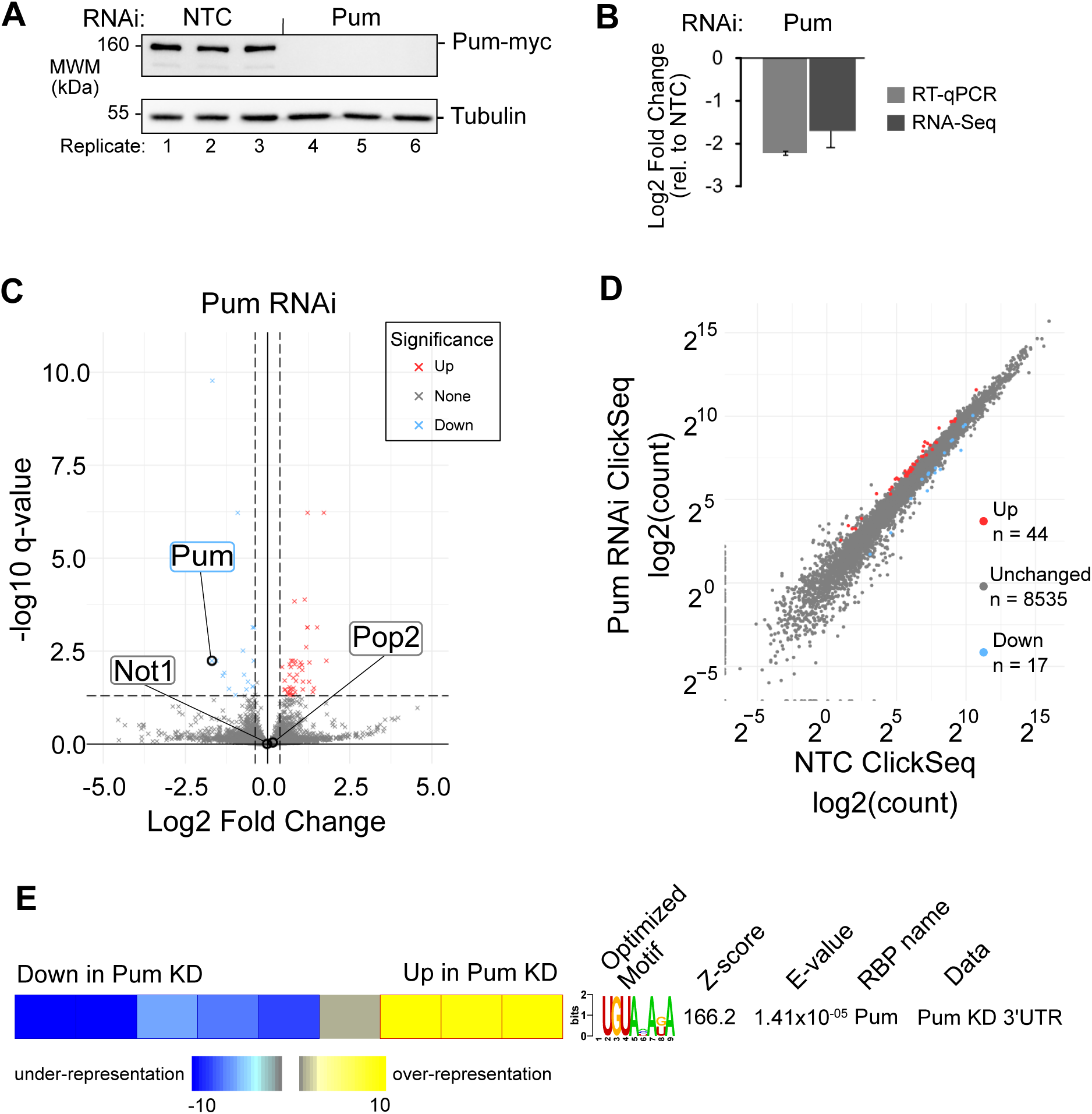
Identification of Pum-regulated transcripts in response to Pum knockdown in *Drosophila* cells. **(A)** Western blot of RNAi knockdown of endogenous myc-tagged Pum protein from 3 biological replicate samples of DL1 Pum-myc cells. RNAi with a non-targeting control (NTC) dsRNA, corresponding to *E. coli lacZ*, served as a negative control. Western blot of tubulin served as a loading control. **(B)** RNAi knockdown of Pum mRNA was confirmed by RT-qPCR and RNA-Seq. Mean log_2_ fold change values +/-standard error of the mean (SEM) are plotted relative to NTC RNAi condition. n=3. **(C)** Volcano plot of statistical significance (q-value) versus mean log_2_ fold change of RNA levels in Pum RNAi relative to NTC RNAi, measured by RNA-seq. Vertical dashed lines indicate a log_2_ fold change value of +/-log_2_(1.3). A statistical significance threshold (q-value ≤ 0.05) is shown with a horizontal dashed line. Red or blue markers (“x”) indicate genes passing both statistical significance and fold change thresholds in positive (Up) or negative (Down) directions, respectively. **(D)** Plot of mean normalized RNA-Seq read counts per kilobase of Pum RNAi versus NTC. Significantly upregulated and downregulated genes are highlighted. **(E)** Identification of RNA-sequence motifs significantly correlated with changes in transcript abundance using FIRE. The distribution of transcript log_2_ fold changes was set in 9 discretized bins, as indicated at the top, and the enrichment or depletion of the identified motif in each bin is shown. Significant enrichment is observed for a motif that is highly identical to the documented Pum binding site, the PRE, in transcripts that are strongly upregulated by knockdown of Pum. The Z-score output by FIRE indicates the information content of the optimized motif. The E-value output by TOMTOM indicates how confidently the motif matched with a known RNA-binding protein.

To measure the effect of Pum on Raf protein expression, the Pum-Myc DL1 cell line was used to create the V5-tagged Raf cell line. The *Raf* gene (FBgn0003079) encodes two transcript variants, both of which code for identical proteins. A single guide RNA plasmid pAc sgRNA Cas9 Raf sg1 was created by inserting annealed primers RJH 327 Raf sg1 Fwd 5′-ttcgTAGATCGCTGTCGCCTTCGG and RJH 328 Raf sg1 Rev 5′-aacCCGAAGGCGACAGCGATCTAc into the BspQ1 site. Note that the 5′ “ttc” or “aac” (in lowercase) are BsqQI cohesive overhangs and the g:c base pair was included as part of the U6 promoter. The V5 tag was integrated onto the N-terminus of the *Raf* coding region, just after the translation initiation codon by co-transfecting the V5-Raf ssODN oligo: 5’-TTCGGGGTCATGGTCACAGCGCATAGTATATAGGATAAAGCAACACC**ATG**<u>ggtaagcctatccctaaccctctcctcggtctcgattctacg</u>TCCAGCGAGTCCAGCACCGAAGGCGACAGCGATCTATACGATC CTTTGGCCGAGGAGCTGCACAACGTCCAGCTCGTCAAACATGTGACCCGCGAGAATATTGATGCC. The V5 tag is indicated by underlined, lower case text and the translation initiation site for Raf in bold text. A homozygous V5-Raf clone was isolated and verified by genotyping, sequencing, and western blot analysis (**Figure 2 and Figure S5**). The following PCR primers that span the exon 3-intron 3 junction, and encompassing the site of V5 tag integration, were used for genotyping and sequencing: RJH323 Raf ex3 Fwd 5′-GCTTGCAAGTGTGTGGG and RJH324 Raf in3 Rev 5′-GGTAGTGTTCAGCTCGGC.

**Figure 2.**
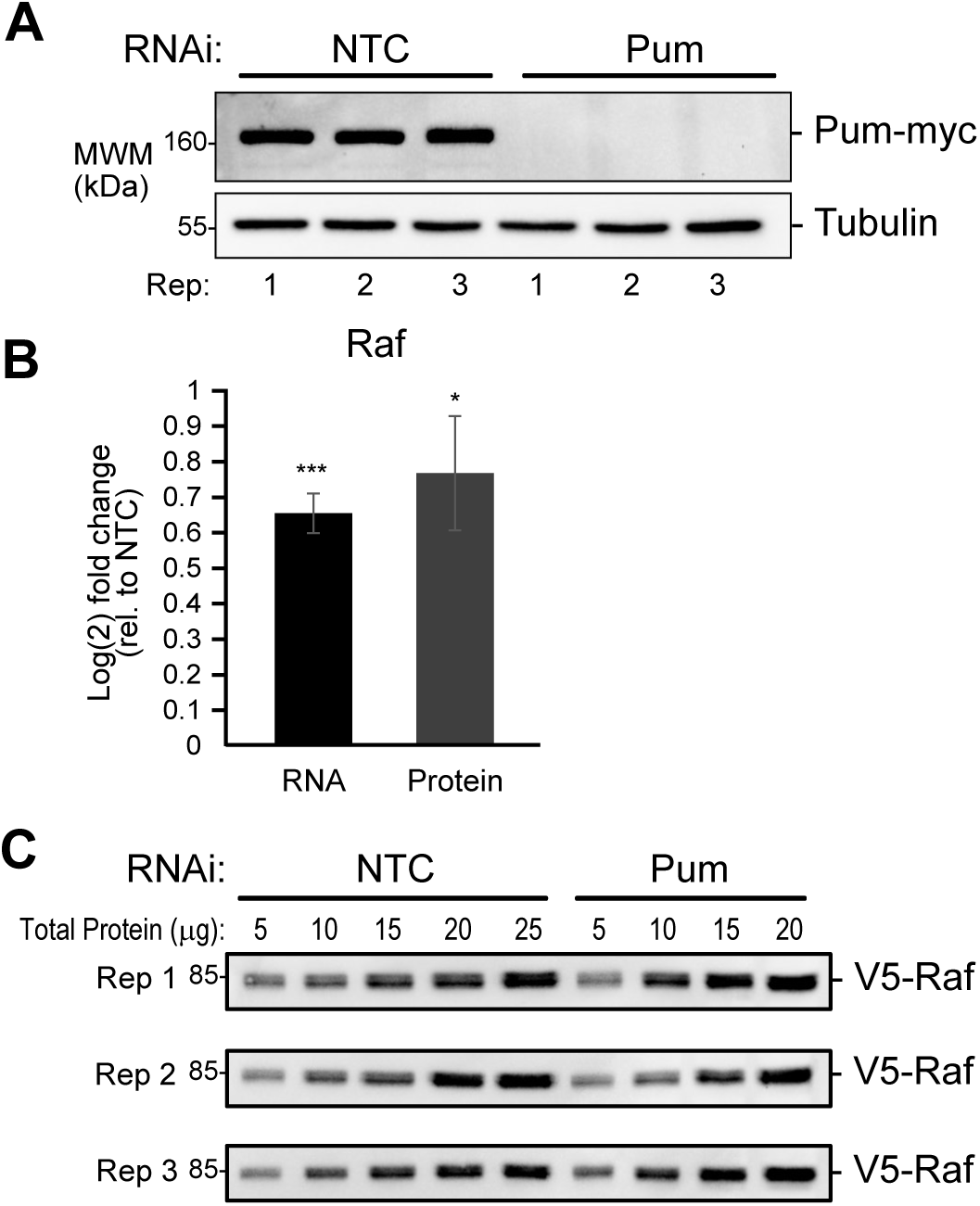
Repression of *Raf* mRNA and protein levels by Pum. **(A)** Western blot confirmed depletion of endogenous myc-tagged Pum by RNAi relative to NTC negative control in 3 biological replicate samples of DL1 V5-Raf, Pum-myc cells. **(B)** Increased expression of Raf mRNA and protein levels in Pum RNAi samples relative to NTC was measured by RT-qPCR and quantitative western blotting, respectively. Mean log_2_ fold change values +/-SEM) are plotted relative to NTC RNAi condition. RT-qPCR measurements were made from 9 replicate samples. Quantitative western blot measurements were made in 3 independent experiments, each of which had 3 biological replicates samples with 4 technical replicate measurements per condition. Significance calling is as follows: * = p<0.05, *** = p<0.001. Details of quantitation are described in the Methods section. **(C)** Representative western blots of V5-tagged endogenous Raf protein in 3 biological replicate samples from an RNAi experiment wherein the effect of Pum RNAi was compared to NTC control in DL1 V5-Raf, Pum-myc cells. The indicated amount of total cellular protein per lane, as measured by DC Lowry assay, for each replicate was analyzed in the western blots.

### Pumilio knockout cell lines

Two homozygous *pumilio* knockout DL1 lines were used in this study. The strategy and tools for generating the CRISPR-Cas9 induced indels and the details of the first clonal line were described in Haugen et al (Haugen et al. 2022). That clonal line (designated Pum KO2 herein) contained a homozygous 10 bp deletion in *pumilio* exon 9, which causes a frameshift after methionine 726 that creates a truncated, nonfunctional 765 amino acid protein that lacks the RNA-binding domain. Here we report an additional homozygous knockout clonal line. This *pumilio* knockout was verified by sequencing a PCR product spanning exon 9 that was amplified from genomic DNA using primers RJH 191 Pum exon 9 fwd primer 5′-AACTGTTTCGCTCGCAGAATCCG and RJH 192 Pum exon 9 rev primer 5′-TGATACGGCTGATTCTCGGCACC (**Figure S3A**). This new *pumilio* knockout line (designated Pum KO1) has a 20 bp deletion of the *pumilio* coding sequence and insertion of 81 additional base pairs, resulting in a net gain of 61 base pairs (**Figure S3B,C**). Therefore, the resulting frameshift after glutamine 725 in this mutant *pumilio* gene adds 8 amino acids followed by a stop codon (**Figure S3D**). This 733 amino acid protein would be nonfunctional due to the absence of the C-terminal RNA binding domain. Additionally, the mRNA produced would be subject to nonsense mediated mRNA decay, consistent with our observation of reduced *pumilio* mRNA in the knockout cells relative to wild type (**Figure S3E**). The RT-qPCR assay and primers for exons 9 and 11 were previously reporter (Haugen et al. 2022).

### RNA Interference

RNAi was performed in Pum-myc tagged DL1 cells. Double-stranded RNA (dsRNA) corresponding to Pum, Not1, and Pop2 were designed using the SnapDragon web-based tool provided by the Harvard *Drosophila* RNAi Screening Center (URL: http://www.flyrnai.org/cgi-bin/RNAi find primers.pl) to minimize potential off-target regions. The effectiveness of these dsRNAs was previously established (Van Etten et al. 2012; Weidmann and Goldstrohm 2012; Arvola et al. 2020). Templates for in vitro transcription were PCR-amplified with primers that add opposing T7 promoters to each DNA strand. The Non-Targeting Control (NTC) dsRNA, corresponding to the E. coli *lacZ* gene, was described previously (Weidman 2012). Primers for generating the Pum, Not1, Pop2, and LacZ dsRNAs were previously described (Weidmann and Goldstrohm 2012; Arvola et al. 2020) including:

T7 LacZ F: 5′-<u>GGATCCTAATACGACTCACTATAGGG</u>TGACGTCTCGTTGCTGCATAAAC

T7 LacZ R: 5′-<u>GGATCCTAATACGACTCACTATAGGG</u>GGCGTTAAAGTTGTTCTGCTTCATC

T7 Pum F: 5′-<u>GGATCCTAATACGACTCACTATAGGG</u>GTCAAGGATCAGAATGGCAATCATGT

T7 Pum R: 5′-<u>GGATCCTAATACGACTCACTATAGGG</u>CTTCTCCAACTTGGCATTGATGTGC

T7 Not1 F: 5’ <u>GGATCCTAATACGACTCACTATAGGG</u>CAAGGACTTCGCCCTGGATG

T7 Not1 R: 5’-<u>GGATCCTAATACGACTCACTATAGGG</u>CATTTGGCTGAGACAAATCCGTCG

T7 Pop2 F: 5’-<u>GGATCCTAATACGACTCACTATAGG</u>GGACACCGAGTTTCCAGGCG

T7 Pop2 R: 5’-<u>GGATCCTAATACGACTCACTATAGGG</u>AAGAAGGCCATGCCCGTCAGC

The dsRNAs were transcribed from these templates using HiScribe T7 high yield RNA synthesis kit (New England Biolabs). The dsRNAs were then treated with DNase and purified using RNA Clean & Concentrator-25 (Zymo Research).

For RNAi experiments, cells (3 x 10^6^ in 1 mL of serum free SDM per well of a 6-well dish) were bathed with 20 µg of dsRNA corresponding to either Pum, Not1, Pop2, or LacZ non-targeting control for 60 minutes. Then 2 mL of SDM containing 5% heat-inactivated FBS was added to each well. After 48 hours, the cells were harvested by centrifugation and suspended in 2 mL Phosphate Buffered Saline (PBS). RNA was purified from 1.5 mL of the cells, and the remaining 0.5 mL was reserved for western blot analysis.

### Western blotting

Cells (∼7.5×10^5^) were harvested by centrifugation at 900 x g for 4 minutes and lysed in radioimmunoprecipitation assay (RIPA) buffer (25 mM Tris-HCl pH 7.6, 1 mM EDTA, 1% NP-40, 1% sodium deoxycholate, 0.1% SDS) containing 2x Complete protease inhibitor cocktail (Roche). Lysates were then cleared of cellular debris by centrifugation at 21,000 x g for 10 minutes. Protein concentration of the supernatant was measured using the detergent compatible (DC Lowry) protein assay kit according to the manufacturer′s directions (Bio-Rad). For each sample, 20 µg of total protein extract was combined with an equal volume of 2 x SDS loading buffer and then heated at 85°C for ten minutes. Samples were then separated by SDS-PAGE and blotted to Immobilon P membranes (Millipore). Membranes were blocked for 1 hour and then the primary antibody (indicated in the respective figure) was applied for one hour at room temperature, or overnight at 4°C on a rocking platform. Antibodies, their dilution factor, and buffer condition are listed below. Membranes were washed three times for ten minutes, then horseradish peroxidase (HRP)-linked secondary antibody was applied for 1 hour at room temperature. After three washes of ten minutes each, chemiluminescent substrate was added to the membrane, which was then imaged with a ChemiDoc Touch instrument (Bio-Rad).

### Antibodies

The following antibodies were used for western blot analysis at the indicated dilutions in either blotto (1x Phosphate Buffered Saline containing 10 mM Na_2_HPO_4_ and 1.8 mM KH_2_PO_4_ at pH 7.4, 137 mM NaCl, 2.7 mM KCl, 0.1% tween-20 and 5% w/v nonfat powdered milk) or TBST (1x Tris-HCl buffered saline containing 50 mM Tris-HCl, pH 7.5, 150 mM NaCl, 0.1% tween-20, and 5% w/v bovine serum albumin), as specified by the antibody′s manufacturer.

Mouse anti-Tubulin (Cell Signaling Technologies, Cat# 3873) at 1:1000 in blotto. Rabbit anti-Myc (Cell Signaling Technologies, Cat# 2278S) at 1:5000 in TBST.

Mouse anti-V5 primary antibody (Invitrogen; catalog no.: R960-25) at 1:5000 dilution in blotto. Goat anti-rabbit-HRP secondary antibody (Cell Signaling Technologies, Cat# 7074P2 or Sigma, Cat# AP187P) at 1:5000 in blotto.

Goat anti-mouse-HRP (Thermo Fisher, Cat# 31430) at 1:5000 in blotto.

#### Quantitative western blotting

To measure the effect of Pum knockdown on endogenous Raf protein levels (**Figure 2**), RNAi of Pum was performed using Pum-myc, V5-Raf DL1 cells. Addition of a V5-tag to Raf was necessary because an antibody to Raf was not available. The cells were plated in 6-well plates at a density of 1×10^6^ cells/mL. To initiate RNAi, cells were bathed with 10 µg of Pum dsRNA, or non-targeting control LacZ dsRNA, for 1 hour in 1 mL of serum-free medium. Thereafter, the medium was exchanged for 2 mL complete serum-containing medium and cells were incubated at 25°C for 90 hours. Cells were then harvested by removing medium and washing the cells with 1xPBS. The cells were lysed in 150 µL RIPA buffer with 2x complete protease inhibitor cocktail (Roche) on ice for 10 minutes, followed by mechanical disruption using a sterile pellet pestle for 20 seconds. Cellular debris was then removed by centrifugation for 10 minutes at 21,000 x g at 4°C. The supernatant was then collected and the total protein concentration was determined by DC Lowry assay.

For quantitation of V5-Raf protein levels, total protein was analyzed by SDS-PAGE for the NTC samples (5, 10, 15, 20, and 25 µg) and Pum RNAi samples (5, 10, 15, and 20 µg), as indicated in the figure. The linear detection range of titration of total protein for the cell extracts was established in optimization experiments (not shown). For each biological replicate, the titrations of total protein were separated on 12% SDS-PAGE gel followed by blotting to Immobilon PVDF membrane (Millipore). Membranes were dried and total protein was stained with Sypro Ruby (Thermo-Fisher) and imaged. Blots were then washed three times in blotto for 1 hour to block and remove the stain. The blots were then probed with anti-V5 antibody overnight at 4°C. The blots were then washed three times with blotto, for 5 minutes per wash, and then were probed with HRP-labeled goat anti-mouse antibody for 1 hour at room temperature. Blots were then washed twice with blotto and then 1xTBS. After incubation with Immobilon ECL substrate (Millipore), the blots were imaged using a ChemiDoc Touch.

Western blot images were analyzed using Image Lab software (Bio-Rad). We performed three independent experiments, each with three biological replicates, and measurements were obtained for 4 amounts of cell extract (5 µg, 10 µg, 15 µg, and 20 µg of total protein), for a total of 36 measurements. For each sample, the measured amount of V5-Raf per band was normalized to total protein in that lane, which was measured prior to sample loading by DC Lowry assay and on the membrane by staining with sypro ruby. We then determined the fold change in normalized V5-Raf protein level in the Pum RNAi condition relative to NTC (Pum KD/NTC) for the same amount of total protein on the same western blot. For example, the V5-Raf band volume for the 5 µg Pum KD sample was compared to the equivalent 5 µg NTC sample on the same blot from the same biological replicate. Mean log_2_ fold change values (Pum KD/NTC) were then plotted with standard error.

### Click-Seq

Click-Seq libraries were generated following the established method (Routh et al. 2015; Routh et al. 2017). RNA was purified using the Maxwell RSC simply RNA tissue extraction kit (Promega) with on-bead DNase I digestion. The RNA concentration was determined using a NanoDrop spectrophotometer. RNA samples were submitted for Agilent Tapestation analysis and RNA integrity number (RIN) values were 8.7-10. Total RNA was poly(A) selected using the NEBNext Poly(A) mRNA Magnetic Isolation Module (New England Biolab).

Reverse transcription was performed in a 20 µL reaction following the manufacturer’s protocol with Superscript III Reverse Transcriptase (Invitrogen) with the following components: 2 µg RNA, 1 µL of 5 mM AzVTP:dNTPs (at a ratio of 1:5 azido-nucleotides AzATP, AzCTP, AzGTP (AzVTP) to dNTPs), 1 µL 50 µM Illumina 6N p7 adapter: 5’-GTGACTGGAGTTCAGACGTGTGCTCTTCCGATCTNNNNNN, 5x Superscript First Strand buffer, DTT, and RNase OUT (Invitrogen). The RNA template was removed after cDNA synthesis by incubating for 20 minutes at 37°C with 10 units RNase H (New England Biolab). Azido-terminated cDNA was purified using a Clean and Concentrator kit (Zymo Research) and eluted in 10 µL of 50 mM HEPES buffer, pH 7.2.

To form the click adapter-linked cDNA, 10 µL of cDNA was incubated for 30 minutes at room temperature with 20 µL DMSO, 3 µL of 5 µM Click-Adapter (5’ Hexynyl-12(N)AGATCGGaaGAGCGTCGTGTAGGGaaAGAGTGTAGATCTCGGTGGTCGCCGTATCATT) and 0.4 µL premixed 50 mM vitamin C and 2 µL of 10mM Cu-TBTA (Lumiprobe). The alkyne-azide cycloaddition of the adapter was catalyzed twice, then the click-linked cDNA was purified with a DNA Clean and Concentrator kit (Zymo Research).

To anneal the remaining Illumina adapters (indexing primer CaaGCAGaaGA CGGCATACGAGATnnnnnnGTGACTGGAGTTCAGACGTGT, where nnnnnn is the index sequence, and universal primer aaTGATACGGCGACCACCGAG), PCR reactions were set up using 2.5 µL each of 5 µM primers, 5 µL click-ligated cDNA, 25 µL 2x OneTaq Standard Buffer Master Mix (New England Biolab) in 50 µL total reactions. Optimized cycling parameters were 94° 4 min; 53° 30 sec; 68°10 min; [94° 30 sec, 53° 30 sec, 68° 2 min] x 20–22 cycles; 68° 5 min. Amplicons were size selected at 200-300 bp on a 2% agarose gel, then gel purified with the Gel DNA recovery kit (Zymo Research). Final yields were determined with a QuBit fluorometer (Thermo-Fisher). Sequencing was performed on pooled samples using single end 75 bp reads on the NextSeq 500 (Illumina) with a high density v2.5 flow cell at the University of Texas Medical Branch Next Generation Sequencing Core Facility.

### Click-Seq data processing

For the Click-Seq samples (Routh et al. 2017; Elrod et al. 2019), sequencing adapters were trimmed and unique molecular identifiers (UMIs) annotated using fastp (version 0.14.1)(Chen et al. 2018), then reads were aligned to the UCSC dm6 genome using hisat2 (version 2.1.0)(Kim et al. 2019). The alignments were then deduplicated using UMI-tools (version 1.0.1) (Smith et al. 2017) and differential expression was analyzed using DESeq2 (version 1.23.110)(Love et al. 2014) and featureCounts (Rsubreads 1.30.9)(Liao et al. 2014). Significance calling was based on adjusted p values (p-adj) and a biological significance cutoff of 1.3-fold change. RNA-Seq data was deposited to the Gene Expression Omnibus as entry GSE##### (In process).

### Identification of PRE-containing genes

FlyBase release 6.38 (February 2021) was used for gene annotations of coding sequences (CDS), 3ʹUTRs, 5ʹUTRs, ncRNAs, miRNAs, and miscRNAs (**Table S1**). Pumilio response element (PRE) processing was performed in R using the seqinR package. Data was then exported to Excel and gene matching based on FBgn, gene name, or gene locus was performed to assign PRE numbers to each gene from our RNA-seq datasets. Transcripts in RNA-Seq data sets that could not be matched to any of the above mentioned FlyBase sequences were discarded; these included but were not limited to genes with withdrawn gene status, pseudogenes, some tRNAs, and mitochondrially encoded genes.

### RIP-Seq

Immunoprecipitation of Pum was carried out as previously described (Laver et al. 2015). The synthetic antibodies, Fab Pum-4 and C1 (Na et al. 2016) were used for Pum-RIP and control-RIP, respectively. Immunoprecipitated RNA and total RNA were extracted using TRIzol (Invitrogen) following the manufacturer’s protocol. The concentration and quality of the RNA samples were checked with Qubit RNA HS Assay Kits (Thermo-Fisher Scientific) and Agilent Bioanalyzer.

Ribosomal RNA was depleted using the following approach. All *Drosophila melanogaster* 5S, 5.8S, 18S, 28S rRNA and 7SL RNA sequences were download from FlyBase (
http://ftp.flybase.net/releases/FB2018_04/dmel_r6.23/fasta/). Each isoform sequence was split into multiple ∼60 nt DNA oligos covering the entire length of its reverse complement strand without overlap. The oligo probes were chemically synthesized by Integrated DNA Technologies Inc. We pooled together equimolar amounts of each of these oligonucleotides to generate oligo probe mixes used for rRNA depletion (**Table S6**).

The protocol for depletion of rRNA and the other RNAs was as published (Adiconis et al. 2013) with minor changes: To deplete target RNAs, we added 1,000 ng of pooled oligos to 1,000 ng of total RNA, incubated the mixture in 1× hybridization buffer (100 mM NaCl, 50 mM Tris-HCl, pH 7.4) in a final volume of 5 μL at 95 °C for 2 min and then slowly lowered the temperature at −0.1°C/s to 50°C. We then added 5 μL preheated RNase H reaction buffer (100 mM NaCl, 50 mM Tris-HCl, pH 7.4, 20 mM MgCl2) with 5 units Hybridase Thermostable RNase H (MACLAB, H39500) to the mix, incubated at 50°C for 30 min and then placed it on ice. We added 10 µL DNase mixture (2x TURBO DNase I Reaction Buffer, 2 units TURBO DNase) to the samples and incubated them at 37°C for 30 min. The samples were cleaned up with 2.2× volume of Ampure XP beads (Beckman Coulter) or Sera-Mag beads for next step library preparation.

RNA-seq libraries were prepared using the Next Ultra II Directional RNA Library Prep Kit (New England Biolab) for Illumina with 96 Unique Dual Index Primer Pairs according to the manufacturer’s protocol. The libraries were sequenced on an Illumina NovaSeq 6000 SP (PE100) at the Next Generation Sequencing Facility, the Centre for Applied Genomics, The Hospital for Sick Children, Toronto, ON, Canada. RIP-Seq data was deposited to the Gene Expression Omnibus as entry GSE240494.

### Reverse transcription and quantitative polymerase chain reaction

RNA was purified from cells using the Maxwell RSC simply RNA tissue extraction kit (Promega) and the concentration was determined using a NanoDrop spectrophotometer (Thermo-Fisher). To confirm Pum knockdown, RT-qPCR analysis of *pumilio* exon 9 and exon 11 in WT and Pum KO DL1 cells was performed according to the method as previously described (Haugen et al. 2022).

A total of 10 µg RNA was taken from each sample for reverse transcription (RT) using GoScript (Promega) reverse transcriptase (5 µg for RT, 5 µg for ‘no RT’ negative control samples). RT reactions were primed with random hexamers and carried out according to the manufacturer’s protocol. cDNA was then diluted with 100 µL water to a final concentration of approximately 41 ng/µL. Quantitative PCR (qPCR) was performed using Go-Taq qPCR Master Mix (Promega) with 2 µL of cDNA or ‘no RT’ sample in a 20 µL reaction volume using 100 or 200 nM final concentration qPCR primers, as indicated for primer sets listed below. Reactions were performed on a Bio-Rad CFX96 instrument using the following cycling parameters: 3 min 98°C, [10 sec 95°C, 30 sec 62/63/64°C, 40 sec 72°C + image] x 39, 60°C-90°C melt curve + image. The fold change induced by RNAi of each mRNA was calculated relative to non-targeting control RNAi condition using the measured Ct values according to the method established by Pfaffl (Pfaffl 2001). Ct values for each target mRNA were normalized to the internal control mRNA encoding ribosomal protein RpL32. Measurements for each target mRNA were repeated in 3 experiments with 3 biological replicates each. Significance calling is as follows: * = p < 0.05, ** = p < 0.01, *** = p < 0.001.

The following primer sets were used to measure *Pop2, Not1, Raf,* and *RpL32* mRNAs.

*Pop2* qPCR primer set produced a 133 bp amplicon 133bp with 100% amplification efficiency (measured according to the method of Pfaffl (Pfaffl 2001)) at 64°C and 200 nM.

RJH510 Pop2 Fwd 5′-AAGTTTAACCTGAGCGAGGACATG RJH511 Pop2 Rev 5′-CAGAGCTCATCAGCAGTTCGG

*Not1* qPCR primer set produced a 157 bp amplicon with 97% efficiency at 64°C and 200 nM.

RJH512 Not1 Fwd 5′-GGACGTGTGCATGGAACTTGATC

RJH513 Not1 Rev 5′-CAGCTGACCTTCCGTGTTTGC

*Raf* qPCR primer set produced a 195 bp amplicon with 100% efficiency at 62°C and 100 nM.

RJH472 Raf Fwd 5′-AGACCTCCTTTGCCGCATCC

RJH473 Raf Rev 5′-GATGCGCGGCCCAATTAAAA

*RpL32* qPCR primer set produced an 85 bp amplicon with 100% efficiency at 62°C and 100 nM.

RC133 RpL32 Fwd 5′-GCCCAAGGGTATCGACAACA

RC134 RpL32 Rev 5′-GCGCTTGTTCGATCCGTAAC

### Determination of dataset overlap significance

P-values for the significance of 2-set or 3-set gene overlaps in the Venn diagrams (**Figures 3C and 4F**) were calculated via one-sided permutation tests, where the gene labels were shuffled (n=1,000) before subsetting based on significance criteria. Fold enrichment was calculated for the genes shared between 2-sets and 3-sets with:

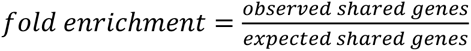

**Figure 3.**
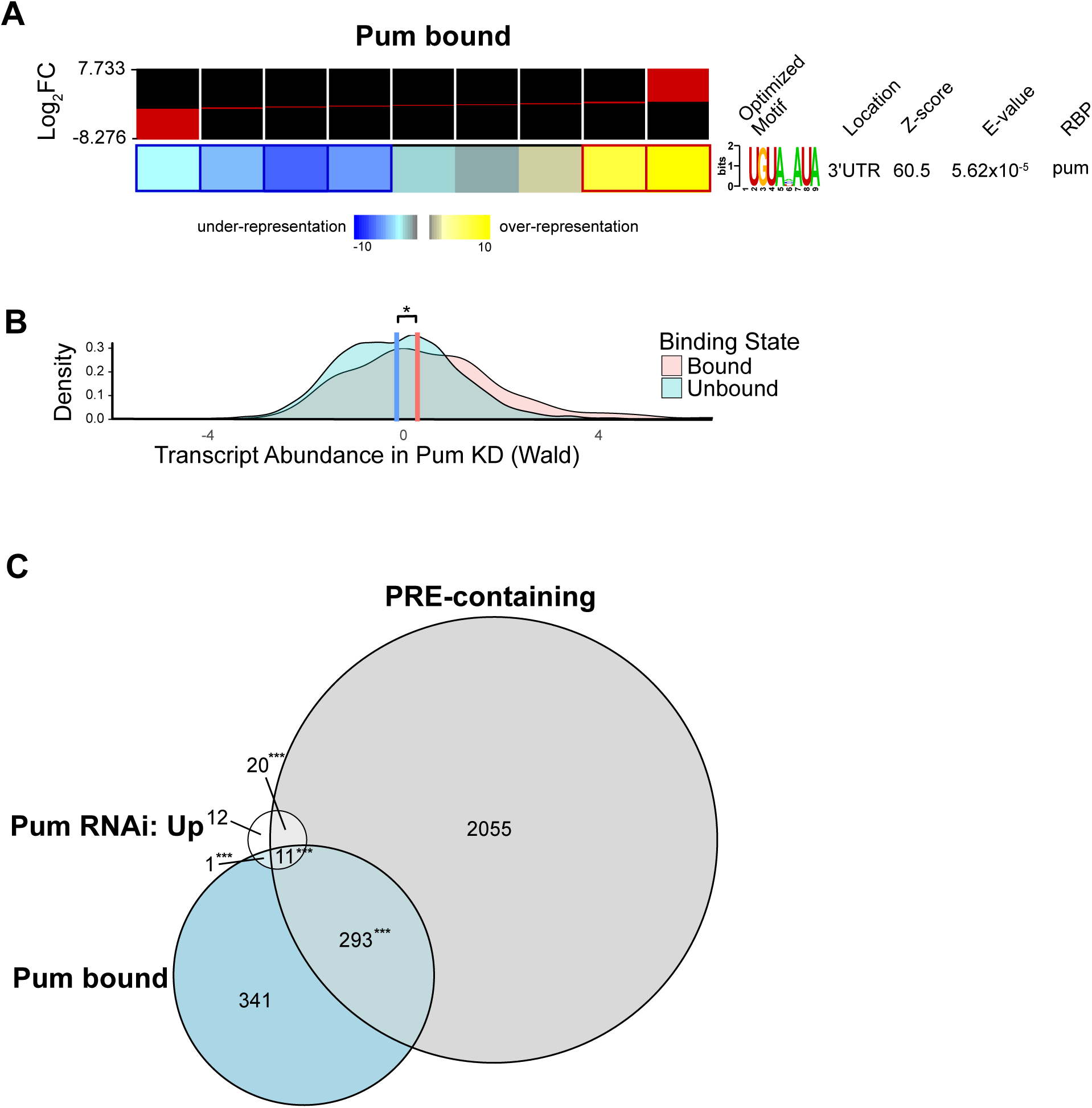
Analysis of Pum-bound mRNAs identifies PRE-containing, Pum repressed direct targets. **(A)** Identification of significantly over-represented RNA-sequence motif in Pum-enriched transcripts from *Drosophila* embryos identified by RIP-Seq and analysis with the FIRE algorithm. The distribution of the log_2_ fold enrichment values for transcripts was set in 9 discretized bins, as indicated at the top, and the over- or under-representation of the identified motif in each bin is shown. Significant over-representation is observed for a motif that is highly identical to the documented Pum binding site, the PRE, in the 3′UTR of transcripts that are strongly enriched in the Pum RIP-Seq. The Z-score output by FIRE indicates the information content of the optimized motif. The E-value output by TOMTOM indicates how confidently the motif matched with a known RNA-binding protein. **(B)** Overlapping density distributions of the RNA abundance changes in response to Pum knockdown, plotted as significance weighted log_2_ fold change (Wald statistic), for genes that were classified as bound (red) or unbound (blue) by Pum RIP-Seq, based on a log_2_ fold change ≥ 1.5 and a q-value ≤ 0.05. **(C)** Venn diagram comparing the overlap of significantly upregulated genes by RNAi of Pum, measured by RNA-Seq, with expressed PRE-containing genes and Pum-bound genes, identified by RIP-Seq. Gene sets are reported in **Table S4**. P-values for the significance of 2-set or 3-set gene overlaps were calculated via one-sided permutation tests, as described in the Methods section. * = p<0.05, ** = p<0.01, *** = p<0.001.

**Figure 4.**
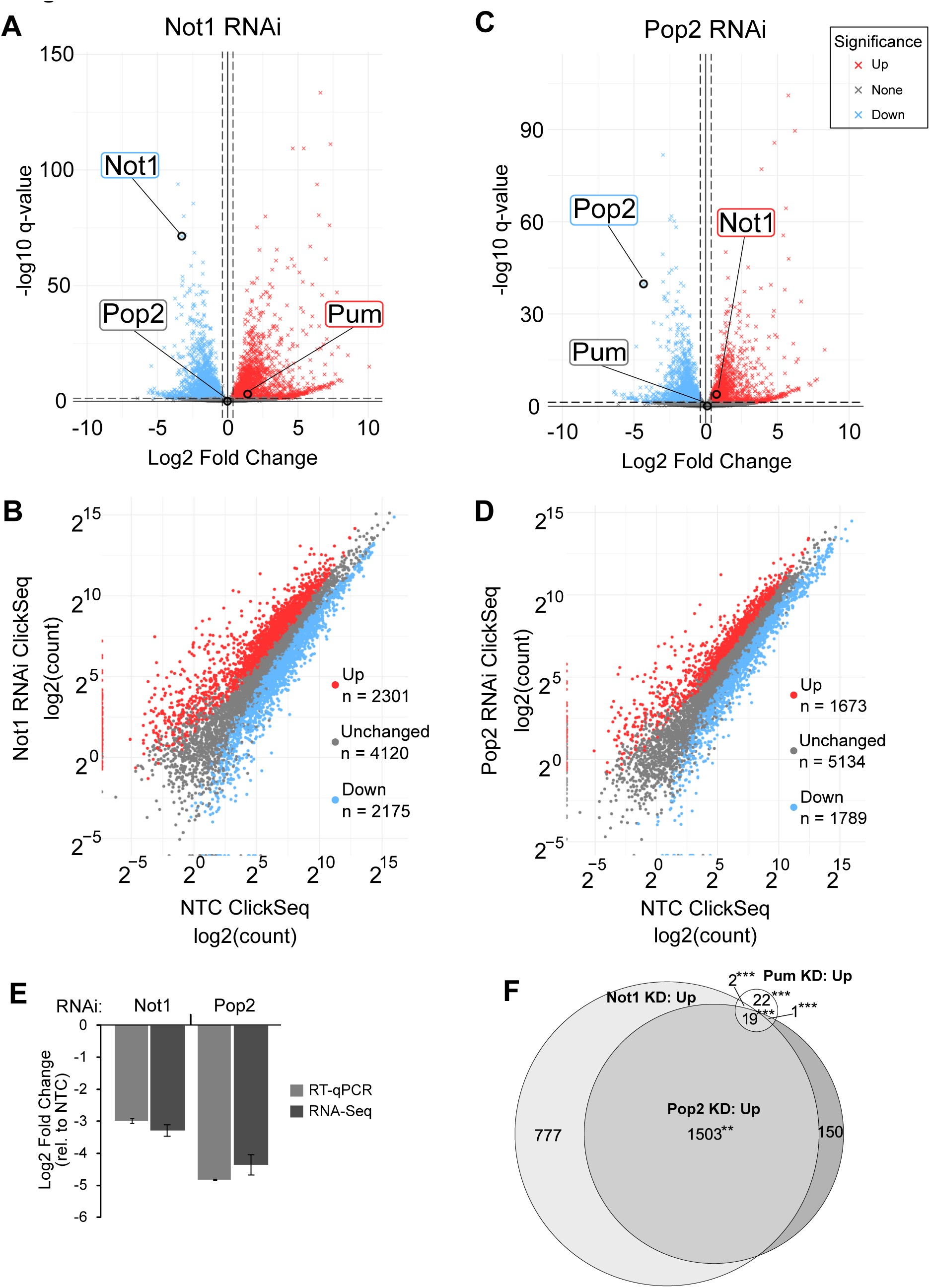
Identification of CCR4-NOT-regulated transcripts in response to knockdown of Not1 or Pop2 in *Drosophila* cells. Volcano plots of RNA level changes measured by RNA-Seq in Not1 **(A)** and Pop2 **(C)** RNAi conditions relative to non-targeting control (NTC) RNAi. The log_2_ fold change (RNAi/NTC) is shown on the x-axis. Vertical dashed lines indicate a log_2_ fold change value of +/-log_2_(1.3). A statistical significance threshold (q-value ≤ 0.05) is shown with a horizontal dashed line. Red or blue markers (“x”) indicate genes passing both statistical significance and fold change thresholds in positive (Up) or negative (Down) directions, respectively. Scatterplots of RNA levels measured in the Not1 **(B)** and Pop2 **(D)** RNAi conditions (y-axis) versus the NTC control (x-axis). **(E)** RNAi knockdown of Not1 and Pop2 mRNA was confirmed by RT-qPCR and RNA-Seq. Mean log_2_ fold change values +/-standard error of the mean (SEM) are plotted relative to NTC RNAi condition. n=3. **(F)** Venn overlap of genes from all 3 KD datasets that were significantly up. Significance was based on the same thresholds as in panels (A-D). All 2-set and 3-set overlaps were significantly enriched (** = p≤0.01, *** = p<0.001, permutation test n=1000), compared to chance (null hypothesis = no correlation between the KDs).

where the expected value was calculated for 2-set overlaps with:

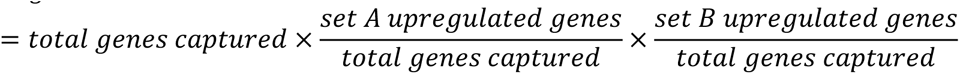

And calculated for 3-set overlaps with the inclusion of a third term:

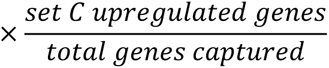

In the case of the overlaps between different experimental types of data, where the total number of genes captured was different for the Click-seq and RIP-seq experiments, the *total genes captured* variable was replaced by the shared intersection of the total genes captured by each experiment.

### Hybrid transcriptome alignment

To check for the presence of Cas9 mRNA reads in the Pum KO1 and KO2 samples, we built a hybrid transcriptome index for kallisto (v0.44.0) (Bray et al. 2016), containing both the FlyBase (Gramates et al. 2022) *D. melanogaster* (r6.38) transcripts along with the sequences for the guide RNAs and mRNAs used in the KO experiments. Each KO sample was run individually, with the arguments `--single --rf-stranded -l 200.0 -s 50.0`. TPM values for reads mapping to each Cas9 mRNA were subsequently pulled from the *abundance.tsv files from kallisto and averaged to calculate an average TPM over the Cas9 mRNA for each KO sample.

### GO-term analysis

GO-term enrichment analysis was performed using iPAGE (Goodarzi et al. 2009) with database files built off of the FlyBase dmel r6.38 genome. For analyses within a single experimental data set (**Figure 5A-C**), genes were discretized into 5 bins based on RNA stability in Pum KD (i.e. `--ebins=5`) and otherwise iPAGE was run with default parameters. For analysis across experimental data sets (**Figure 5D**), genes were discretized into 4 categories based on whether they shared the same directionality in all three RNAi knockdown datasets. The “Down in all 3” and “Up in all 3” categories were based on whether the genes met our significance criteria (log_2_ fold change ≥ +/-log_2_(1.3), q-value ≤ 0.05) in either direction. Genes belonging to the “Neutral in all 3” category, consistently did not meet our significance criteria in any of the 3 KD data sets. Genes that had any other combination of directionalities across data sets were added to the “Different directions” category. iPAGE was run with the argument `--exptype=discrete` and otherwise with default parameters.

**Figure 5.**
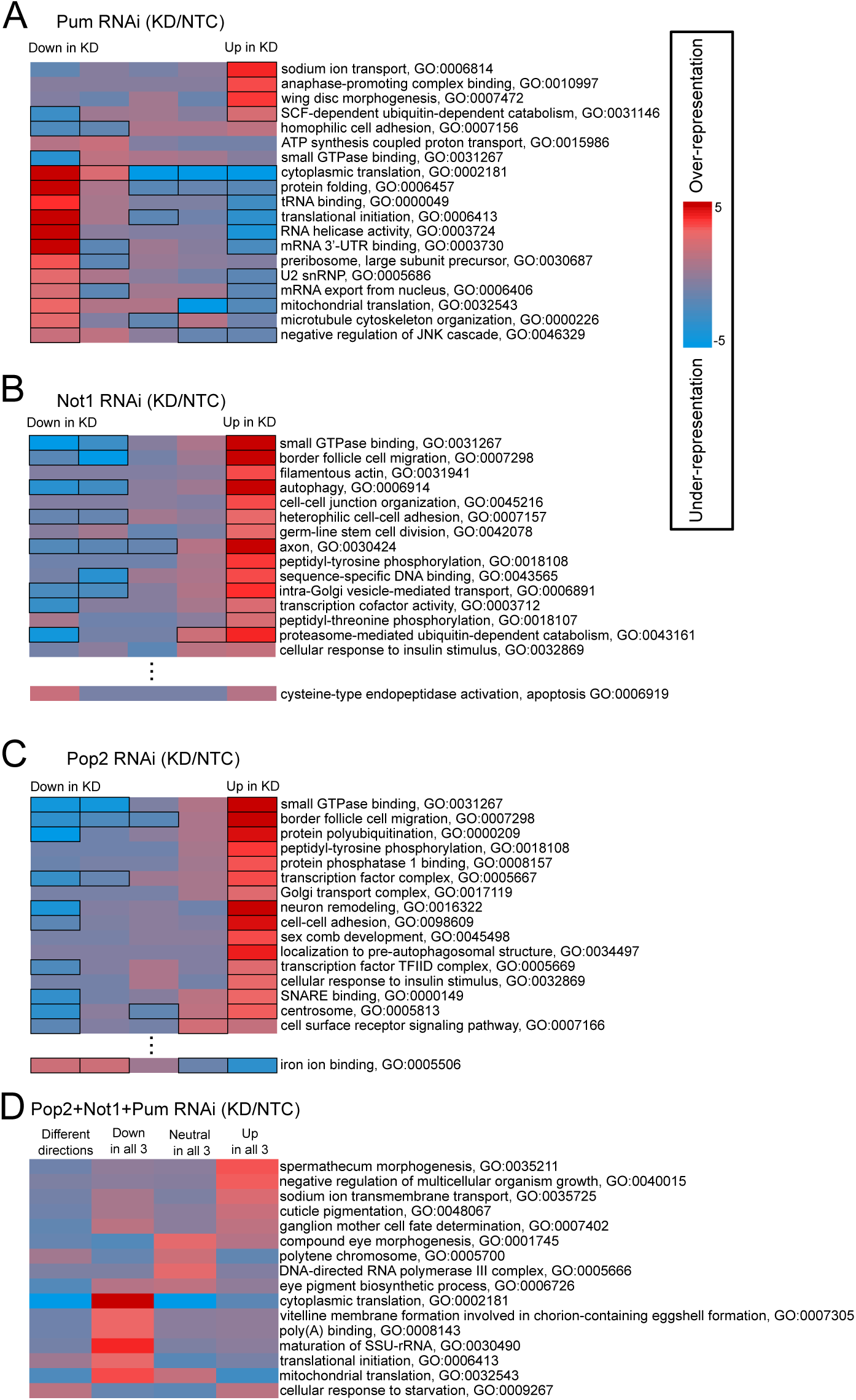
Gene ontology enrichment analysis of differentially expressed transcripts in response to Pum and CCR4-NOT knockdown relative to negative control RNAi (NTC). **(A)** GO terms showing significant mutual information with the observed log_2_ fold changes upon Pum knockdown, measured in our RNA-seq dataset. The log_2_ fold changes were discretized into 5 equally populated bins, ranging from downregulated on the left (Down in KD) to upregulated on the right (Up in KD). GO terms showing significant mutual information, calculated using iPAGE, are listed on the right. The color of each cell shows the magnitude of the P-value for significance of the enrichment (red) or depletion (blue) at that cell, scale is shown on the right-hand side of the figure; black bordered cells individually show P-values ≤ 0.05 after Bonferroni correction across their row. The plotting and highlighting conventions described here apply to all panels of the figure except for panel D, which is a discrete analysis and lacks the indication of Bonferroni corrected P-values ≤ 0.05. **(B)** As in panel A, for Not1 knockdown data. **(C)** As in panel A, for Pop2 knockdown data. **(D)** GO term enrichment analysis using iPAGE of four categories of genes based on their downregulation (Down in all 3) or upregulation (Up in all 3) upon knockdown of Pum, Not1, and Pop2. The “neutral” category encompasses genes that were not differentially expressed in any knockdown. The “Different directions” category includes the genes that responded in opposition between any two knockdown conditions.

### Naive motif discovery and analysis

We identified motifs that were informative of the observed gene regulatory changes in our Click-seq expression data, from each of the Pum, Not1, and Pop2 knockdowns, using a development version of Finding Informative Regulator Elements (FIRE)(Elemento et al. 2007; Goodarzi et al. 2009). We used FIRE to evaluate both the 5’UTR and 3’UTR regions for all three datasets. The FIRE-discovered motifs were then cross-referenced against the CISBP-RNA database of known RNA-binding protein (RBP) motifs in *D. melanogaster* using the TOMTOM motif comparison tool to quantify the similarity of our discovered motifs to the motifs already known (Gupta et al. 2007). Motifs were considered to be both informative and a good match to known RBPs if they had a Z-score ≥ 50 from FIRE and an e-value ≤ 0.05 from TOMTOM.

### Modeling gene regulation

The following features were generated as previously described (Wolfe 2020). 1) The number of perfect PRE matches present. 2) The maximum number of PREs clustered together. 3) The AU-content surrounding the PRE site(s). 4) The absolute distance of the PRE into the 3′UTR. 5) The relative location of the PRE in the 3′UTR. 6) Percent codon optimality (data was obtained from previously reported values in *Drosophila* (Burow et al. 2018). 7) Pum binding data from RIP-seq data in the form of log_2_ fold enrichment values. 8) the length of the 3′UTR. These features were used as the independent variables for our dependent variable, which was the Wald statistic observed from the Pum KD Click-Seq experiment.

Linear regression models were fitted to the Pum KD expression data using the lm() function from the stats (v4.2.0) package in R (v4.2.3; R Core Team 2023)(https://www.R-project.org/). Generalized additive models were fitted using the gam() function from the mgcv v1.8-40 with the method argument set to “REML” (Wood 2011). The importance of individual features was evaluated with the Bayesian information criterion (BIC) applied to leave-one-out variations of the linear or generalized additive model. The BIC was calculated using the BIC() function also from the R stats (v4.2.0) package (Wood 2011).

## RESULTS

### RNA-Seq identifies new Pumilio-regulated mRNAs

To identify mRNAs that are regulated by Pum, we knocked down its expression in the embryo-derived *Drosophila* Line 1 (DL1) cells using RNAi (Pum KD) and measured the impact on RNA levels using RNA-Seq. We reasoned that while not a complete loss of function, this transient depletion of Pum is likely to be less prone to adaptive changes in gene expression due to the speed with which it can be accomplished. We performed RNA-Seq on libraries generated from poly(A) selected RNAs using the Click-Seq method (Routh et al. 2015). Differential expression in the Pum KD condition was determined relative to the non-targeting control (NTC) RNAi using dsRNA corresponding to the *E. coli LacZ* gene (Weidmann and Goldstrohm 2012; Arvola et al. 2020; Haugen et al. 2022). Three biological replicates were analyzed for each condition. For this experiment, DL1 cells were modified using CRISPR/Cas9 genome editing to introduce a myc epitope tag into the *pumilio* gene (**Figure S1**), which enabled confirmation of depletion of Pum protein by western blotting (**Figure 1A**). Knockdown of *pumilio* mRNA was also confirmed by RT-qPCR and RNA-Seq (**Figure 1B**). The observed level of Pum depletion is consistent with previous studies that demonstrated stabilization of PRE-containing reporter mRNAs and alleviation of Pum-mediated repression (Weidmann et al. 2014; Arvola et al. 2020; Haugen et al. 2022).

The levels of more than 8600 genes were measured in this experiment. The results are summarized in **Table 1**, with full data reported in **Table S2**. Differentially expressed mRNAs were identified using a ≥ 1.3-fold change threshold and, as a cutoff for statistical significance, an FDR adjusted p-value of ≤ 0.05 (**Figure 1C, 1D**), as previously established (Van Etten et al. 2012; Weidmann and Goldstrohm 2012; Bohn et al. 2018; Enwerem et al. 2021). By these criteria, 44 genes were significantly upregulated in the Pum KD condition. For example, the aquaporin big brain (*bib*), which is involved in neurogenesis and modulates notch signaling, was among the most highly upregulated genes (> 3.2-fold, p adj = 0.005). In contrast, only 17 genes were significantly decreased by Pum KD. Given the documented role of Pum in repression of gene expression (Arvola et al. 2017), hereon we primarily focus our analysis on the upregulated genes. Considerations and limitations of this RNAi approach are addressed in the Discussion.

**Table 1.** Significantly affected genes by Pum, Not1, and Pop2 knockdown and Pum knockout. The number of genes that were significantly up or down regulated in each knockdown and knockout experiment are reported. Significance was determined based on whether the gene had a q-value ≤ 0.05 and a +/-fold change ≥ 1.3. Complete datasets are listed in **Table S2** for Pum KD and **Table S3** for Pum KO.

| RNAi Knockdowns |  |  |
| --- | --- | --- |
|  | Up | Down |
| Pum KD | 44 | 17 |
| Not1 KD | 2301 | 2175 |
| Pop2 KD | 1673 | 1789 |
| Overlap of Not1 + Pop2 KD | 1522 | 1523 |
| Overlap of Not1 + Pop2 + Pum KD | 19 | 10 |
| CRISPR-Cas9 Knockouts |  |  |
| Pum KO1 | 1776 | 2173 |
| Pum KO2 | 1573 | 1878 |

### Enrichment of PRE motifs in Pumilio-regulated mRNAs

If the mRNAs that are upregulated by Pum KD are direct targets, then they should contain one or more binding sites for Pum. Using the well-characterized RNA-binding specificity of Pum (5′-UGUANAUA, which constitutes a Pumilio recognition/response element, or PRE)(Arvola et al. 2017), we cataloged the PRE-containing genes in the *Drosophila* genome, documenting the number and location of PREs within the annotated transcripts. **Table 2** provides an overview and the complete list is reported in **Table S1**.

**Table 2.**
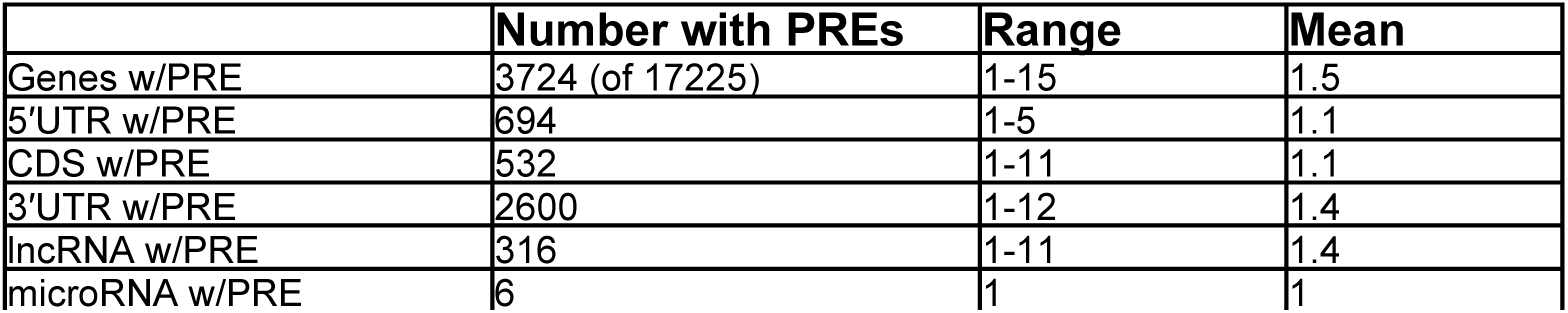
Summary of PRE-containing genes in *Drosophila melanogaster*. The number of genes containing Pumilio Response Element sequences is listed, along with the range of PREs and the overall mean. Unique genes with one or more mRNA isoforms that contain a PRE are included in the total number. The complete dataset is reported in **Table S1**.

We found that 31 of the 44 genes upregulated ≥ 1.3 by Pum KD contain one or more PREs (**Table 3)**. These genes encode proteins involved in neurodevelopment, germ cell development, transposon suppression, metabolism, cell proliferation, and signaling (i.e., Notch, Wnt, TGF-beta, and MAPK/ERK pathways). Of these genes, only the *Raf* oncogene was previously connected to Pum, based on reporter gene assays showing that the *Raf* 3′UTR confers repression by Pum (Haugen et al. 2022). The location and number of PREs in each Pum-repressed mRNA is shown in **Table 3**. It is notable that there is a statistically significant enrichment in the intersection between genes that are Pum-regulated and those containing a PRE in the 3′UTR (**Table 4**). Indeed, most of the genes that were upregulated in response to Pum KD have PREs located in the 3′UTR (29 of 31). The remaining two genes upregulated by Pum KD (*CBP, Jheh2*) have a single PRE in their 5′UTR – and none elsewhere in the transcript – suggesting the potential for Pum repression via 5′UTR binding sites, although an indirect effect on these two transcripts remains possible.

**Table 3.** Drosophila genes that are significantly upregulated in response to Pum RNAi and contain one or more PRE. The flybase gene name and symbol for each gene are listed, along with a summary of its function and the location of PREs within the transcript. For genes with multiple mRNA isoforms, PRE parameters are based on the longest isoform. Genes that were also upregulated in both Pum KO cell lines are indicated in bold text. Genes with transcripts bound by Pum in embryos, identified by RIP-Seq, are indicated with a “+”.

| Pum-repressed, PRE-containing Gene |  |  | PRE location |  |  |  |
| --- | --- | --- | --- | --- | --- | --- |
| Flybase ID | Symbol | Summary | 5'UTR | CDS | 3'UTR | Bound |
| **FBgn0000077** | **amx** | neurodevelopment, Notch pathway | 0 | 0 | 2 | + |
| FBgn0001128 | gpdh1 | glycerol-3-phosphate dehydrogenase 1, metabolism | 0 | 0 | 2 | + |
| **FBgn0002973** | **numb** | neurodevelopment, Notch pathway | 0 | 0 | 1 | + |
| FBgn0011704 | RnrS | deoxynucleotide biosynthesis, ribonucleoside diphosphate reductase, small subunit | 0 | 0 | 1 | + |
| **FBgn0024889** | **Kap-alpha1** | spermatogenesis, nuclear import, karyopherin | 0 | 0 | 1 | + |
| FBgn0024994 | Ugalt | UDP-Galactose transporter | 0 | 0 | 3 | + |
| FBgn0028360 | cdc7 | promotes DNA replication, protein kinase | 0 | 0 | 2 | + |
| **FBgn0031491** | **alpha4GT1** | α1,4-galactosyltransferase 1, glycosphingolipid biosynthesis | 0 | 0 | 1 | + |
| **FBgn0036502** | **CG7841** | function and phenotype unknown | 0 | 0 | 2 | + |
| **FBgn0037110** | **ORMDL** | negative regulator of sphingolipid synthesis | 0 | 0 | 2 | + |
| FBgn0039135 | CG13603 | function and phenotype unknown | 0 | 0 | 1 | + |
| **FBgn0000180** | **bib** | neurodevelopment, Notch pathway | 0 | 1 | 1 |  |
| **FBgn0003079** | **Raf** | oncogene, serine-threonine kinase | 0 | 0 | 2 |  |
| **FBgn0003716** | **tkv** | germ cell development, TGF beta receptor for dpp signaling | 0 | 0 | 3 |  |
| **FBgn0010473** | **tutl** | neurodevelopment, axon guidance, motor control, IgG superfamily transmembrane protein | 0 | 0 | 1 |  |
| FBgn0011288 | snap25 | SNARE complex, neurotransmitter release | 0 | 1 | 2 |  |
| FBgn0015766 | Msr-110 | integral membrane component, molecular function unknown, expressed in digestive system, embryo germ band, prothoracic gland | 0 | 0 | 1 |  |
| **FBgn0024245** | **dnt** | muscle development, cell morphogenesis, differentiation, Ryk family of Wnt receptor tyrosine kinase | 0 | 0 | 1 |  |
| **FBgn0025626** | **CG4281** | function unknown, expressed in gonad and digestive system | 0 | 0 | 2 |  |
| FBgn0026144 | CBP | sarcoplasmic calcium binding protein | 1 | 0 | 0 |  |
| FBgn0031401 | papi | piRNA pathway, repression of transposable elements during meiosis | 0 | 0 | 1 |  |
| **FBgn0034405** | **Jheh2** | juvenile hormone epoxide hydrolase 2 | 1 | 0 | 0 |  |
| **FBgn0034611** | **MFS16** | integral membrane component, molecular function unknown | 0 | 0 | 2 |  |
| FBgn0035696 | Best2 | predicted chloride transport channel | 0 | 1 | 2 |  |
| FBgn0038460 | CG18622 | function and phenotype unknown | 0 | 0 | 4 |  |
| **FBgn0039045** | **Ctns** | amino acid transmembrane transport, Cystine/H<sup>+</sup> symporter | 0 | 0 | 1 |  |
| FBgn0040334 | Tsp3A | Notch signaling, tetraspanin superfamily transmembrane protein | 0 | 0 | 1 |  |
| FBgn0040348 | CG3703 | GTPase activator, expressed in embryonic brain and larval ventral nerve cord | 0 | 0 | 1 |  |
| FBgn0041629 | Hexo2 | beta-N-acetylglucosaminidase involved in carbohydrate metabolism | 0 | 0 | 1 |  |
| **FBgn0264090** | **CG43759** | function unknown | 0 | 0 | 3 |  |
| FBgn0283509 | Phm | neuropeptide biosynthesis, peptidylglycine-α-hydroxylating monooxygenase | 0 | 0 | 2 | |

**Table 4.** Functional and statistical relationship of Pum-regulated, bound, and PRE-containing genes. Untranslated region: UTR; Coding sequence: CDS; Fold Enrichment: FE; p-values from permutation test, n=1000.

| PRE Location: |  | 5'UTR | CDS | 3'UTR |
| --- | --- | --- | --- | --- |
| Pum-regulated & PRE-containing | p-value | 0.778 | 0.171 | < 0.001 |
|  | log<sub>2</sub>(FE) | -0.403 | 1.105 | 1.953 |
| Pum-regulated & Pum-bound | p-value | < 0.001 | < 0.001 | < 0.001 |
|  | log<sub>2</sub>(FE) | 1.943 | 1.977 | 1.956 |
| PRE-containing & Pum-bound | p-value | < 0.001 | 0.013 | < 0.001 |
|  | log<sub>2</sub>(FE) | 1.220 | 0.605 | 1.249 |
| Pum-regulated, PRE-containing, & Pum-bound | p-value |  |  | < 0.001 |
|  | log<sub>2</sub>(FE) |  |  | 0.575 |

We sought to identify cis-RNA elements associated with the observed changes in gene expression caused by Pum KD in the RNA-Seq dataset. To do so, we applied the motif discovery tool Finding Informative Regulatory Elements (FIRE), which evaluates the significance (assessed via mutual information) of the presence or absence of a motif while making as few prior assumptions as possible (Elemento et al. 2007). We used FIRE to search for motifs in the 3′ and 5′UTR regions (Goodarzi et al. 2009). To determine if these motifs correspond to known binding sites of RBPs, we used the tool TOMTOM (Gupta et al. 2007) to cross reference the motifs discovered by FIRE to the CISBP-RNA database, which catalogs known RBP motifs, including for *Drosophila* RBPs (Ray et al. 2013). Motifs were considered to be both significantly informative for the gene expression and a confident match to known RBP motifs if they had a Z-score ≥ 50 from FIRE and an E-value ≤ 0.05 from TOMTOM, respectively. The results revealed that an over-represented motif in the 3′UTR of genes upregulated by Pum KD significantly matches the RNA-binding specificity of Pum (**Figure 1E**)(Gerber et al. 2006; Weidmann et al. 2016; Arvola et al. 2017; Wolfe et al. 2020). We noted that several additional motifs were identified by this analysis (**Figure S2**), though their recognition by RBPs and functional relationship to Pum are obscure. Taken together, our results identify multiple new Pum-regulated target genes and emphasize the importance of 3′UTR PREs in Pum-mediated repression.

### Differential gene expression analysis in Pumilio knockout cells

As an additional approach to identify Pum-regulated target mRNAs, we used CRISPR-Cas9 to inactivate the *pumilio* gene (Haugen et al. 2022). Two Pum knockout (KO) DL1 cell lines were clonally isolated and genotyped. Pum KO1 contains a 20 bp deletion and an 81 bp insertion after glutamine 725 that appends 8 additional amino acids and a premature stop codon (**Figure S3A-D**). Pum KO2 has a 10 bp deletion that creates a frameshift and premature stop codon after methionine 726, as we recently reported (Haugen et al. 2022). Both Pum KO1 and KO2 proteins lack an RNA-binding domain, thereby rendering them non-functional. Both Pum knockouts also produce significantly less mRNA, as measured by RT-qPCR (**Figure S3E** and (Haugen et al. 2022)) and RNA-Seq (**Figure S4A, B**), consistent with their propensity to be degraded by the nonsense mediated decay. We then performed RNA-Seq on three replicates of each Pum KO line and compared differential gene expression relative to parental wild type (WT) DL1 cells. RNA levels from 9924 genes were measured. The complete loss of function of Pum was anticipated to result in upregulation of a greater number of transcripts and, indeed, many more genes were significantly affected compared to the Pum KD experiments (**Figure S4**, summarized in **Table 1**, with complete dataset reported in **Table S3**). Overall, 1247 genes were significantly upregulated ≥ 1.3-fold in both the Pum KO1 and KO2 lines (**Table S3**).

We anticipated that the upregulated genes would be enriched for direct Pum targets in a manner consistent with Pum-mediated repression. Of the genes with significantly increased expression in both Pum KOs, 399 contain PREs (**Table S3**), including documented Pum targets *hunchback* (*hb*) and *brat* (Wharton and Struhl 1991; Murata and Wharton 1995; Harris et al. 2011; Arvola et al. 2017). Though its expression in wild type DL1 cells is low, *hunchback* was among the most highly upregulated, increasing by 8-to 14-fold in Pum KO lines (p adj < 0.05). When we examined the overlap of significantly upregulated genes captured by both KO and KD approaches, we found 16 PRE-containing genes, including *Raf, bib, numb*, and *tutl* (**Table 3**). These mRNAs encode factors involved in germ cell, muscle, and neurodevelopment and key signaling pathways. These observations provide corroborating evidence for Pum-mediated repression of a novel collection of direct target mRNAs.

While the preceding results from the Pum KOs are informative, we also found reason for cautious interpretation. First, the significantly affected genes in the KOs were skewed towards down regulation (45% up vs. 55% down), contrary to the expected effect of loss of Pum activity. This contrasts with the results of transient Pum KD, wherein the majority of affected genes were upregulated (72% up vs 28% down). Second, the majority of differentially expressed genes in the Pum KOs (68%) do not appear to be direct Pum targets in that they lack PREs, again unlike the case of the transient knockdown. Third, contrary to expectation, the PRE motif is not significantly over-represented in the genes that are upregulated by knockout of Pum. Instead, our de novo motif enrichment analysis using FIRE and TOMTOM identified over-represented several motifs, including those corresponding to Pabp2 and B52/SRp55 RNA-binding proteins, both of which participate in mRNA processing and regulation (**Figure S2**) (Roth et al. 1991; Mayeda et al. 1992; Ring and Lis 1994; Benoit et al. 1999). However, the patterns of their motif enrichment did not correspond to upregulation by Pum KO and therefore likely result from indirect or secondary effects.

Our observations led us to suspect that constitutive loss of Pum and/or a technical issue may have contributed to the changes in gene expression. Several caveats are worth consideration. Pum is essential for viability in animals and, though we succeeded in knocking out Pum in the cultured cells, adaptive changes in gene expression may have occurred to compensate for the loss of Pum function. Further, Cas9 may have persisted in the Pum KO cells with the potential to cause off-target effects. Indeed, examination of the RNA-Seq data for the presence of Cas9 mRNA indicated that the Pum KO1 and KO2 lines had average TPMs of 22.5 and 13.4, respectively, placing it in the ∼75-percentile of expressed genes. Deeper inspection of transcriptomes revealed higher expression of DNA damage response genes (e.g., CG7457/FBgn0035812, a human TONSL ortholog) in the Pum KO lines, potentially reflecting DNA damage by Cas9. We note that the CRISPR-Cas9 KOs used here were generated using standard procedures in the field (see Methods); we suggest that evaluation of residual Cas9 expression may be prudent in other contexts. As a result of these considerations, we chose to primarily focus our analysis on the RNAi dataset, which did not show signs of competing or adaptive effects.

### Expression of the Raf oncogene is regulated by Pumilio

Based on the observed upregulation of *Raf* mRNA in Pum KD and KO RNA-Seq datasets, we performed further validation. *Raf* encodes a serine-threonine kinase which activates the MAPK/ERK pathway to regulate proliferation and differentiation (Hayashi and Ogura 2020). The *Raf* gene expresses two mRNA isoforms, *Raf-RA* and *Raf-RE*, which are produced by separate promoters and differ solely in their 5′UTRs. Both *Raf* mRNAs share the same protein coding sequence and 460 nt 3′UTR, which contains 2 PREs. To validate Pum regulation, we measured changes in *Raf* mRNA and protein levels in response to Pum KD. As a *Drosophila* Raf antibody is not available, we engineered a V5 epitope tag on the N-terminus of the *Raf* coding sequence using CRISPR-Cas9 in the DL1 cells wherein the *pumilio* gene was myc-tagged. We clonally isolated a homozygous V5-Raf cell line, which was confirmed by genotyping, sequencing, and western blotting (**Figure 2 and S5**). The effect of Pum knockdown on *Raf* mRNA and protein was then measured relative to a non-targeting control RNAi. To assess reproducibility, these measurements were made in 3 independent experiments, each with 3 biological replicates. Depletion of Pum-myc protein was confirmed by western blot (**Figure 2A**). To measure changes in *Raf* mRNA, we performed RT-qPCR on total cellular RNA using optimized primers that detect both mRNA isoforms. Consistent with the RNA-Seq results, Pum KD significantly increased *Raf* mRNA abundance (**Figure 2B**). The magnitude of the increase of *Raf* mRNA determined by RT-qPCR (fold change = 1.7, p < 0.001) closely matched the RNA-Seq data (fold change = 1.6, p adj = 0.029). In parallel, we performed quantitative western blotting on titrated amounts of cell extracts to measure changes in V5-tagged Raf protein between Pum KD and NTC conditions (see Methods for details). Pum KD significantly increased Raf protein to the same degree as *Raf* mRNA level (**Figure 2B, C**). Pum regulation of Raf is further supported by our recently published reporter gene data showing that the *Raf* 3′UTR mediates repression by Pum dependent on its 2 PREs (Haugen et al. 2022). Together with the RNA-Seq Pum KD and KO data, these results support Pum-mediated repression of Raf expression. In the Discussion, we relate this important finding to the biological roles of Pum and Raf in *Drosophila* development.

### Integration of Pum-bound, PRE-containing, Pum-regulated mRNAs

As an additional parameter to characterize direct regulation of target genes, we incorporated new experimental evidence for Pum-binding to mRNAs in the context of early embryos, chosen because Pum has documented regulatory roles in this context (Lehmann and Nusslein-Volhard 1987; Arvola et al. 2017). To do so, Pum was immunoprecipitated using a synthetic monoclonal Fab antibody (Laver et al. 2015) and Pum-associated mRNAs were identified by RNA-Seq (RIP-Seq). Transcript enrichment in the Pum RIP-Seq was determined relative to a negative control immunoprecipitate, with a fold-enrichment cutoff of > 1.5 fold and FDR corrected p-value < 0.05. Of the 7034 expressed genes, 695 transcripts from 646 unique genes were significantly enriched by Pum (**Table S4**).

We then performed de novo motif enrichment analysis using FIRE. In this application, genes were stratified into equivalent sized bins based on their log_2_ fold enrichment. FIRE identified a motif that is significantly over-represented in Pum bound transcripts and matches the PRE consensus (**Figure 3A**). Indeed, a total of 329 of the Pum-bound genes contain PREs (**Table S4**). The Pum transcript itself was among the most highly enriched, consistent with a previous report of autoregulation (Laver et al. 2015). FIRE analysis detected significant over-representation of the PRE motif in the 3’UTR of Pum-bound transcripts (**Figure 3A**) but not in the 5’UTR or CDS. Further analysis of all PRE containing and Pum-bound transcripts indicates that Pum is significantly more likely to bind mRNAs that have a PRE, regardless of where that PRE is located, though there is a slight enrichment bias towards 3’UTR PREs, followed by 5’UTR PREs, over CDS PREs (**Table 4**).

We investigated whether evidence of Pum binding to transcripts was informative in relation to increased transcript abundance in the Pum KD RNA-Seq data. Based on the RIP-Seq data, RNA-binding by Pum had a statistically significant association with increased RNA abundance caused by Pum KD (**Figure 3B**, **Table 4**). Thus, Pum binding to a transcript is informative of Pum-mediated repression of its abundance, consistent with direct regulation.

We then sought to identify the highest confidence direct Pum target RNAs by integrating three categories of evidence: 1) upregulation in response to Pum KD, 2) Pum binding from RIP-Seq, and 3) the presence of one or more PRE. Of the 44 genes that were significantly upregulated by Pum KD, 31 have PREs and 12 genes were detected in Pum-bound transcripts (**Figure 3C** and **Table S4**). A total of 11 transcripts matched all 3 criteria, including *amx, numb,* and *cdc7* (**Table 3**).

### Concordant regulatory effects of Pumilio and CCR4-NOT

We previously showed that the CCR4-NOT complex is necessary for Pum repression of PRE-containing reporter genes; specifically, RNAi of Pop2 or Not1 subunits reduced Pum:PRE mediated repression and mRNA degradation (Arvola et al. 2020). Pop2 is one of two deadenylase enzymes in the CCR4-NOT complex, and is thought to provide the major cytoplasmic deadenylase activity in *Drosophila* (Temme et al. 2010; Temme et al. 2014). Not1 is the structural backbone upon which the CCR4-NOT complex assembles. Further, multiple repression domains of Pum directly bind to Not1, 2 and 3 subunits of CCR4-NOT (Arvola et al. 2020; Haugen et al. 2022). Importantly, the effect of the Pum:CCR4-NOT mechanism on natural target mRNAs remained unknown. If Pum uses CCR4-NOT to repress most mRNAs, then depletion of Pop2 or Not1 should increase the levels of Pum target mRNAs. In addition to its role in mRNA decay by Pum, CCR4-NOT participates in other regulatory processes, and therefore we anticipated that depletion of Not1 or Pop2 would stabilize a broader spectrum of transcripts (Chicoine et al. 2007; Eulalio et al. 2009; Temme et al. 2014; Raisch and Valkov 2022).

To measure the impact of CCR4-NOT on the transcriptome, Pop2 and Not1 were knocked down by RNAi using previously established conditions (Arvola et al. 2020). RNA-Seq was performed on three replicates for each RNAi condition and compared to the control (NTC). Notably, this experiment was conducted in parallel with the Pum KD described in **Figure 1**. Knockdown of Not1 and Pop2 led to increased expression of 2301 and 1673 genes (**Figure 4A-D**, **Table 1, and Table S2**), in a manner consistent with the role of CCR4-NOT in RNA decay. Knockdown of Not1 and Pop2 was confirmed by RT-qPCR and RNA-Seq (**Figure 4E**).

We then assessed the overlap of mRNAs stabilized by Pum, Not1, and Pop2 depletion. A total of 8,596 genes were measured across all three datasets. We found that Not1 KD and Pop2 KD had a statistically significant overlap of 1522 upregulated genes, consistent with their mutual function in the CCR4-NOT complex (**Figure 4F**, **Table 1, Table S5**). Comparison of Pum, Not1, and Pop2 KD datasets revealed a statistically significant overlap of 19 significantly upregulated genes (**Table S5**), including high confidence Pum targets listed in **Table 3** (*bib, numb, tkv, amx, Tsp3A, Raf, cdc7, RnrS, CG13603,* and *CBP*). Together, these datasets reveal new PRE-containing natural transcripts that are negatively regulated by Pum and CCR4-NOT.

### Gene ontology enrichment analysis of Pum and CCR4-NOT datasets

To gain insight into the network of genes affected by Pumilio, we performed gene ontology (GO) term enrichment analysis on our RNA-Seq data using iPAGE (Goodarzi et al. 2009). iPAGE allows the identification of annotations (GO terms) that have significant mutual information with expression profiles or other quantitative data. Key advantages of this mutual information approach include: 1) that any correlation structure can be detected (even if it is not monotonic; e.g., enrichments specifically among unchanged genes, or among both up- and down-regulated genes) and 2) that redundant terms can be removed in a principled way by requiring each new GO term to add significant new information (assessed based on the conditional mutual information), thus avoiding over-calling of related GO terms. Transcripts were separated into 5 bins based on their significance-weighted log_2_ fold change in response to Pum KD. GO terms that were significantly over- or under-represented in those categories were identified, as indicated by the heatmap (**Figure 5A**).

The results reveal significantly affected molecular and biological processes and pathways modulated by Pumilio. Genes upregulated by Pum KD were significantly over-represented with GO terms including sodium ion transport, wing disc morphogenesis, and cell adhesion (**Figure 5A**). In addition, genes that mediate proteolysis to regulate the cell cycle were also over-represented, including the anaphase-promoting complex and Skp1-cullin 1-F-box (SCF) ubiquitin ligase complex. In the Discussion, we relate these observations to Pum’s known phenotypes and biological functions. In contrast, RNA-related processes, such as cytoplasmic translation and RNA helicase activity, were over-represented in the downregulated category.

We also analyzed the RNA-Seq datasets from Not1 KD (**Figure 5B**) and Pop2 KD (**Figure 5C**). A diverse group of GO terms was over-represented in upregulated gene sets, many of which are identical or related between Pop2 KD and Not1 KD, consistent with their coordinated function in the CCR4-NOT complex. Among those in common are signaling pathways, post-translational modifications, cell adhesion and migration, and intracellular trafficking. GO terms over-represented in the downregulated genes included glycolytic process, cytoplasmic translation and rRNA processing (see **Figures S6-S7** for expanded lists).

We also performed a comparative GO term enrichment analysis of the Pum KD, Not1 KD, and Pop2 KD gene sets. Genes were binned into four categories based on their responses in the three conditions (**Figure 5D**): 1) upregulated genes in Pum and Not1 and Pop2 knockdowns (Up in all 3), 2) genes that did not change in any of the knockdowns (Neutral in all 3), 3) genes that decreased in all knockdowns (Down in all 3), and 4) genes that responded in different directions among the three knockdowns (Different directions). Interestingly, the ‘Up in all 3’ category has enriched categories involved in morphogenesis of spermathecum and compound eye, negative regulation of growth, sodium ion transmembrane transport, and ganglion mother cell fate determination. RNA polymerase III transcription was over-represented in the ‘Neutral in all 3’ category, along with eye pigmentation and polytene chromosome terms. In contrast, categories related to translation were over-represented in the ‘Down in all 3’ gene list. Taken together, these results provide new insights into the overlapping and unique regulatory networks of Pum and CCR4-NOT. Future research in vivo will be important to pursue these new relationships and their potential phenotypic consequences.

### Functional association of PRE features with Pum-mediated repression

To further our understanding of Pum repression, we investigated contextual features of Pum-responsive mRNAs. We restricted our analysis to the Click-Seq RNAi data to ensure that we were building the models on the most robust experimental data. We initially focused on the number of PRE sites, adenine and uracil (AU) content, and the relative location of PRE sites within the 3′UTR, which we previously found were strong predictors of binding and transcript destabilization by human Pumilio orthologs in human cells (Wolfe et al. 2020). We began by investigating the relationship of these features in the 3′UTRs to the Pum RNAi KD differential expression results in *Drosophila* cells. Each feature was assessed in relation to the significance-weighted log_2_ fold change (Wald statistic) in transcript abundance.

First, we examined the relationship of the number of PREs to regulation by Pum. The abundance of transcripts with up to 3 PREs significantly increased in response to Pum KD (**Figure 6A**), consistent with direct repression by Pum. Interestingly, transcripts with 4-9 PREs were associated with a decrease in RNA abundance, and their median abundance dipped below the median of transcripts with no PREs, but the differences at higher PRE counts were not statistically significant. We noted that the number of transcripts with 4 or more PREs comprised less than 0.7% of all transcripts considered, compared to 21.1% of transcripts with one or more PRE sites. Since low numbers reduce the power of the Wilcoxon rank sum test, we repeated the test to compare transcripts with zero perfect PRE sites to all transcripts that had one or more PRE present in the 3′UTR. In this case, we found that the overall presence of one or more PRE sites led to a significant increase in RNA abundance upon Pum depletion (**Figure 6B**).

**Figure 6.**
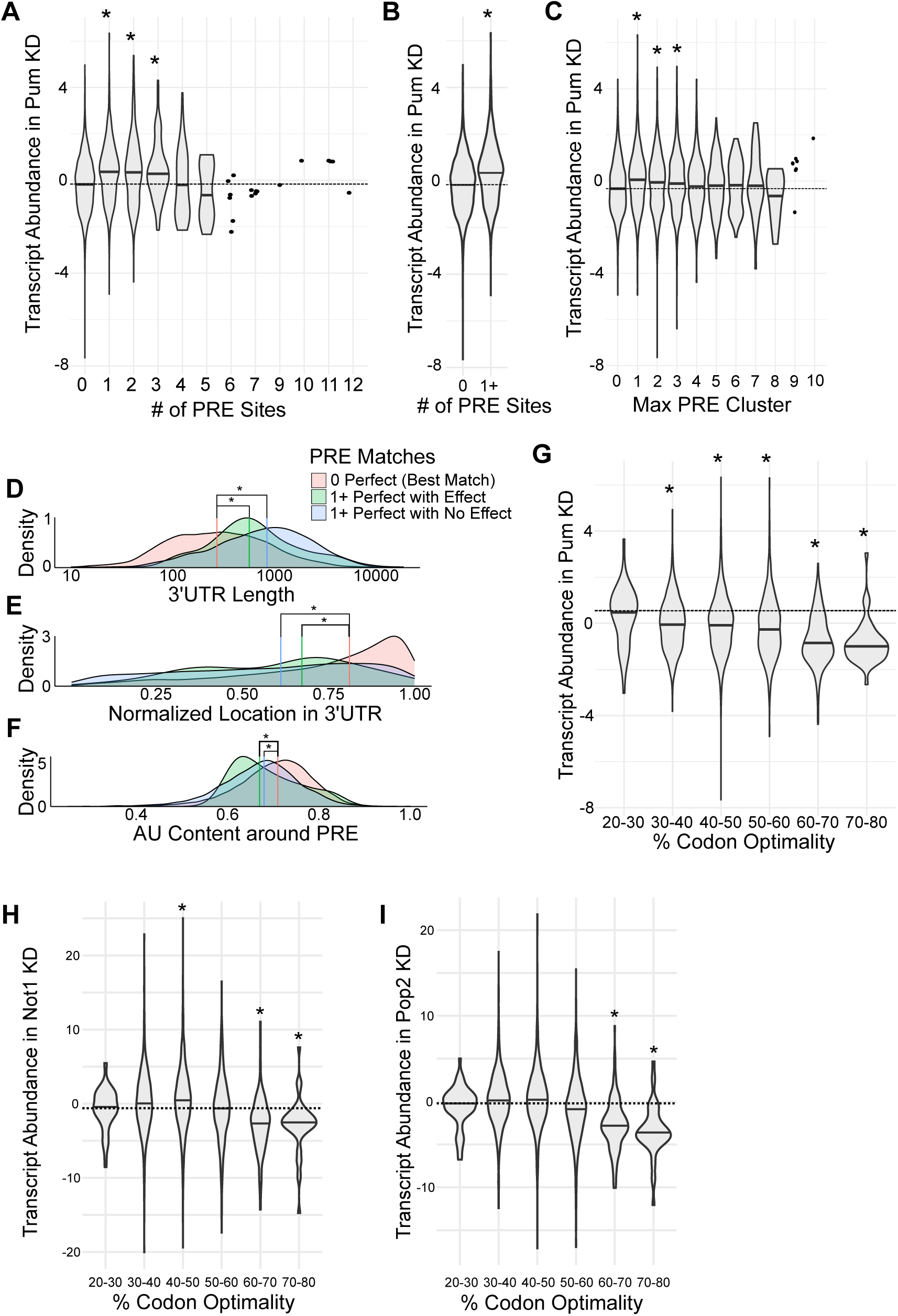
Functional associations between PRE features, codon optimality, and differential gene expression in response to knockdown of Pum, Not1, and Pop2. **(A-C)** Density distributions of transcript abundance in Pum KD for transcripts **(A)** with different numbers of 3′UTR PREs sites, **(B)** with or without a PRE site in the 3′UTR, and **(C)** with different the maximum numbers of 3′UTR PRE sites clustered within a 100bp window. Density medians are indicated with black lines within each violin. The null median is shown as a dashed line across the width of each panel. Statistical significance (p-value ≤ 0.05, Wilcoxon rank sum test) is indicated by an asterisk above each violin. **(D-F)** Overlapping density distributions of **(D)** 3′UTR length, **(E)** the normalized location of a 3′UTR PRE, and **(F)** the %AU content within a 100bp window of 3′UTR PREs, for genes that had no perfect PREs (red), genes that had at least one PRE and passed the significance thresholds (LFC +/-log2(1.3) and q-value ≤ 0.05) (green), and genes that had at least one PRE but did not pass the significance thresholds (blue). Density medians are indicated by their corresponding color-coded vertical line. Statistical significance (p-value ≤ 0.05, Wilcoxon rank sum test) is indicated by an asterisk. **(G)** As in panels (A-C), for transcripts binned based on their content of optimal codons. **(H)** As in panel G, for Not1 knockdown data. **(I)** As in panel G, for Pop2 knockdown data.

We next examined the potential functional relationship of clustering of PREs in 3′UTRs. PRE clustering was assessed within a sliding window of 100 nts. The maximum number of clustered PREs showed a statistically significant increase in the median RNA abundance in response to Pum KD up to 3 clustered PREs, but not when there were 4 or more clustered PREs (**Figure 6C**).

We examined three additional sequence-context features of interest: 3′UTR length, the length-normalized location of PRE sites, and the percent AU sequence content surrounding PRE sites. For this analysis, we divided transcripts into 3 groups defined in **Table 5**. We then tested whether there were differences in the median values of the three groups for each feature. The PRE-containing mRNAs (groups 2 and 3) had significantly different median values for 3′UTR length (**Figure 6D**), PRE location (**Figure 6E**), and AU content (**Figure 6F**), relative to the median value of these features for mRNAs without a PRE (group 1). The results show that PREs tend to occur in longer 3′UTRs (**Figure 6D**), toward the 3′ end (**Figure 6E**), and in a context with slightly less AU content (**Figure 6F**). However, between the groups that contained PREs and either were significantly impacted by Pum depletion (group 2) or were not (group 3), there was no significant difference in median values for any of the three sequence-context features enumerated above. Therefore, this analysis did not detect a statistically significant role of 3′UTR length, PRE location, or AU content as determinants of Pum-mediated repression in *Drosophila* cells.

**Table 5:**
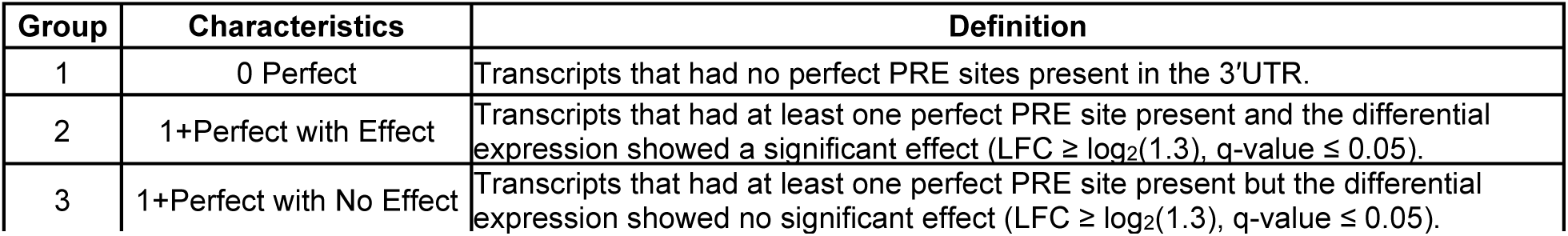
Transcript group criteria for feature evaluation. Defining criteria for the transcript groups 1, 2, and 3. Groups were defined based on whether or not the 3′UTR of the transcript contained a PRE site that perfectly matches the consensus sequence 5′-UGUANAUA and whether or not the transcript appeared significantly repressed by Pum.

### Codon optimality is a functional determinant of Pum repression

Codon optimality is a parameter that relates synonymous codon usage to the efficiency of translation and degradation of mRNAs in eukaryotic organisms (Bae and Coller 2022; Wu and Bazzini 2023). We investigated the potential relationship of codon optimality to Pum-mediated repression. For this purpose, we used previously established *Drosophila* codon stabilization coefficients (Burow et al. 2018). Transcripts were binned based on their overall percentage of optimal codons. We then analyzed the relationship of percent optimal codons to the significance-weighted log_2_ fold change (Wald statistic) in transcript abundance in response to Pum KD. Surprisingly, codon optimality showed a significant linear correlation with Pum-induced changes in RNA abundance (**Figure 6G**). Transcripts with low percentage optimal codons had increased abundance in response to Pum depletion, whereas transcripts with higher percentage optimal codons were less responsive. The most straightforward interpretation of this correlation is that transcripts with lower codon optimality are more susceptible to degradation by Pum than those with higher codon optimality.

As described in the Introduction, based on research in budding yeast, the CCR4-NOT complex is proposed to mediate codon optimality-mediated mRNA decay (Buschauer et al. 2020; Bae and Coller 2022). However, the relationship of CCR4-NOT-mediated degradation with codon optimality in higher eukaryotes in this process remains to be determined. We interrogated the functional relationship of percent codon optimality to the change in transcript abundance caused by knockdown of Not1 and Pop2. The results revealed that transcripts with high codon optimality are significantly less likely to be affected by either Not1 or Pop2 depletion. Collectively, our results indicate that the inherent translation efficiency and/or stability of the mRNA can modulate its responsiveness to Pum and CCR4-NOT.

### Computational modeling of Pum repression

We applied linear regression modeling to further assess which features contribute the most to Pum-mediated repression of mRNA levels. We utilized a leave-one-out analysis to determine the relative importance of the following 8 features: 1) Pum binding, 2) PRE count, 3) 3′UTR length, 4) max clustered PREs, 5) AU content, 6) percent codon optimality, 7) PRE location, and 8) normalized PRE location. The Bayesian information criterion (BIC) was used to compare the performance of the models. The first 6 features were found to be informative, and the BIC for the 6-feature model was slightly better than the model using all 8 features, with an approximate 11-point drop. In both models, the percent variance in the expression data explained by the input features was between 11-12%, indicating that one or more unknown features contributing to Pum-mediated repression are missing from the model. We also tested a more complex generalized additive model (GAM) with the same 6 features and found that it produced a less informative model, thus we focused on the linear modeling. The informative features for the Pum KD data, in descending order, were: Pum binding (RIP-seq log_2_ fold enrichment) values (ΔBIC = 112.57), AU content around the PRE site (ΔBIC = 80.77), codon optimality (ΔBIC = 53.09), number of perfect PRE sites (ΔBIC = 43.61), the number of PRE sites clustered together (ΔBIC = 27.22), and 3′UTR length (ΔBIC = 18.19) **(Fig 7)**. The top feature, Pum binding, had roughly 140% the impact on the 6-feature model BIC than that of the next highest impact feature. Taken together, the results of our analysis indicate that Pum-binding and sequence-context features are determinants of whether a transcript responds to Pum. Importantly, our full 6-feature model accounts for a low percentage of data variation (adjusted R-squared = 0.115); therefore, it is likely that additional functional determinants remain to be discovered.

**Figure 7.**
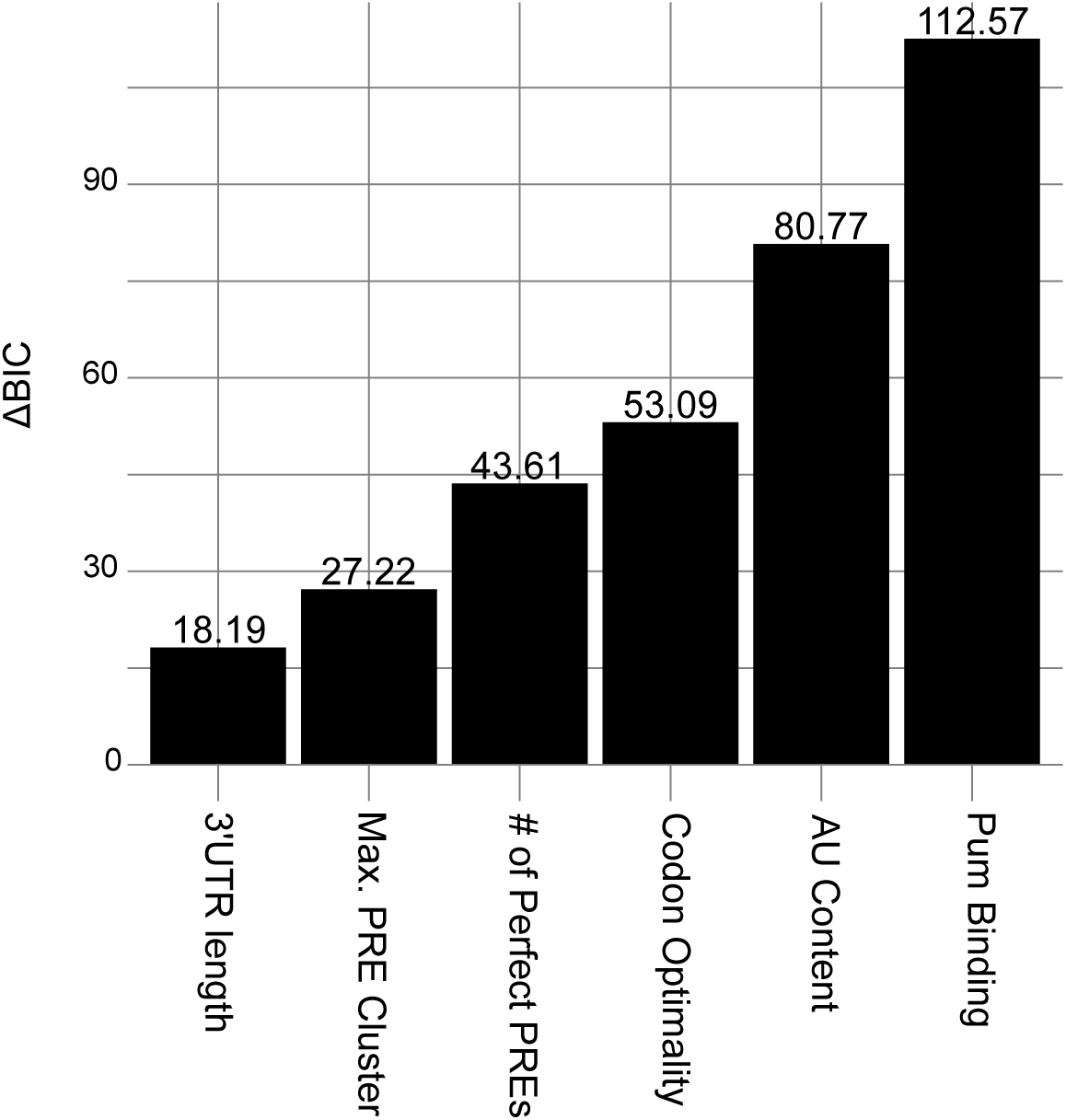
Computational modeling of Pum-mediated regulation of mRNAs identifies informative features. Leave-one-out analysis of a linear regression model applied to 6 influential features of Pum expression, labeled on the x-axis. The information contributed to the model by each feature was quantified with the change in the Bayesian Information Criterion (ΔBIC), where the difference was calculated as BIC_1 left out_ - BIC_full model_ resulting in a positive ΔBIC value for informative features in the model.

## DISCUSSION

*Drosophila* Pumilio has long served as an archetype of an RNA-binding regulator with crucial biological functions (Arvola et al. 2017). This study identifies new transcripts that are upregulated in response to Pum depletion and deletion. By integrating multiple criteria, we winnowed that list to define high confidence, functionally repressed, direct Pum target mRNAs. The functions of proteins encoded by these Pum repressed target genes provide important insights into the regulatory roles of Pum (**Table 2**). We found that Pum represses genes involved in neural, muscle, and germ cell development. Pum repressed targets also function in wing morphogenesis, cellular proliferation, and differentiation. Components of the DNA replication (e.g. *cdc7, RnrS*), lipid (e.g. *Gpdh1*) and sphingolipid metabolism (e.g. *α4GT1, ORMDL*), and transposable element suppression pathways (e.g. *papi*) are directly repressed by Pum. Key components of signaling pathways are repressed by Pum including Wnt (e.g. *dnt*), TGF-beta (e.g. *tkv*), and MAPK/ERK (e.g *Raf*). Strikingly, 4 Notch signaling components are direct Pum targets (*amx, bib, numb, Tsp3A*). A previous study provides additional evidence for Pum-mediated repression of four targets (*bib*, *RnrS*, *Kap-alpha1*, and *CG4281*) in embryonic neural progenitors (Burow et al. 2015). Future work should explore the functional role of Pum regulation of the high-confidence targets in vivo, in the context of development and physiology. Intriguingly, some of the same Pum-repressed signaling pathways identified in our study (Wnt, TGF-beta, MAPK) are targeted by Pum orthologs in other species (Bohn et al. 2018; Goldstrohm 2018). For example, human and *C. elegans* Pumilio orthologs regulate MAPK/ERK signaling in stem cells (Lee et al. 2007).

Pum repression of *Raf* is relevant to their shared role in wing morphogenesis. Pum was previously shown to negatively affect MAPK/ERK signaling by epidermal growth factor receptor to control wing development (Kim et al. 2012). Our results establish that *Raf* is a high-confidence direct target of Pum repression, providing a means by which Pum can modulate MAPK-dependent wing development. Regulation of *Raf* by Pumilio proteins may be conserved in mammals, which have three *Raf* homologs (ARAF, BRAF, RAF1). The ARAF mRNA has 2 PREs in its 3′UTR and was reported to be bound by Pumilio proteins in both human and mouse, whereas RAF1 and BRAF do not have PREs, though the association of BRAF mRNA with human PUMs has been reported (Galgano et al. 2008; Zhang et al. 2017; Bohn et al. 2018). While our approach successfully identified new directly regulated Pum targets, the list of significantly repressed RNAs is modest compared to the breadth of transcripts that contain PRE(s) or are bound by Pum in embryos. We consider several relevant factors. First, Pum may repress some mRNAs by additional mechanisms, such as inhibition of translation in lieu of mRNA decay (Weidmann et al. 2014; Arvola et al. 2017). Second, the endogenous level of Pum in wild type cultured cells may be at a limiting threshold, in which case mRNAs could escape repression unless the level of active Pum became elevated. In support of this possibility, we previously showed that over-expression of Pum strengthened repression of target mRNAs in a dose-dependent manner (Weidmann and Goldstrohm 2012; Arvola et al. 2020; Haugen et al. 2022). Third, Pum activity may be modulated by a factor(s) that is absent from our experimental system. For example, the Nanos RNA-binding protein can promote Pum-mediated repression but is expressed at an exceedingly low level in DL1 cells (**Table S2,** *nos*)(Weidmann et al. 2016; Arvola et al. 2017). Fourth, we considered that residual Pum in the knockdown condition might be sufficient to maintain repression of some RNAs. To address this issue, we created Pum knockout cell lines and measured differential gene expression relative to wild type cells. As anticipated, substantially more genes were significantly differentially expressed in the Pum knockout cells, and while many of those contain PREs (including bona fide direct Pum targets like *hunchback* and *Raf*), the PRE was not over-represented in the upregulated transcripts. We suspect that adaptive changes in gene expression may have occurred during clonal isolation of the Pum knockout cells that buffer the loss of Pum activity. We also found evidence of residual Cas9 expression in these cells, which could contribute to off-target effects (Guo et al. 2023). For future analysis of regulated mRNA decay, a rapid, efficient, and transient means of depletion of a regulatory protein would be ideal, such as conditional degrons.

Our data provide insight into mRNA features that contribute to repression by Pumilio. Foremost, the location, number, and density of PREs in a transcript are functionally important. In particular, PREs located in the 3′UTR are significantly correlated with repression by Pum, as is the number and density of PREs up to 3 sites. Notably, we also observed functional relationships between the location, number, and density of PREs to repression by human Pumilio proteins (Bohn et al. 2018; Wolfe et al. 2020). Our computational modeling of Pum repression emphasizes six important functional determinants. Three relate to the Pum:RNA interaction, including evidence of Pum binding, number of perfect PRE sites, the number of clustered PRE sites, and PRE location. The fourth, length of 3′UTR, may relate to the propensity of longer UTRs to contain regulatory sequences and structures. The fifth, surrounding AU content of PREs, may reflect the reduced stability of AU base pairs relative to GC rich sequences. Such AU content may facilitate accessibility of functional PREs. The sixth determinant, percent codon optimality, is discussed below. The modeling also indicates that additional determinants remain to be identified to accurately predict the regulation of a gene by Pum without functional data. This remains a universal challenge for the gene regulation field. We speculate that collaborative activities of other RNA-binding factors or RNA structural features, which remain to be discovered, may help specify targets of Pum-mediated repression.

Pum repression is mediated by direct recruitment of the CCR4-NOT deadenylase complex (Arvola et al. 2020; Haugen et al. 2022). Prior to this work, the Pum:CCR4-NOT repression mechanism was established using reporter mRNAs and supported by a single example of a natural target mRNA in ovaries, *mei-P26* (Joly et al. 2013; Weidmann et al. 2014; Arvola et al. 2020). Our results extend the Pum:CCR4-NOT mechanism to a group of natural target mRNAs in *Drosophila* cells. Interestingly, some of the Pum repressed mRNAs we identified were not affected by depletion of the Pop2 and Not1 CCR4-NOT components. Residual CCR4-NOT in the RNAi conditions may have been sufficient to support Pum activity. Alternatively, we speculate that Pum may repress these transcripts by another mechanism, such as decapping-mediated mRNA decay and/or antagonism of poly(A) binding protein (Weidmann et al. 2014; Arvola et al. 2020).

Our results also emphasize the broader effect of CCR4-NOT on the transcriptome, consistent with the growing list of pathways and RNA-binding regulatory proteins that utilize CCR4-NOT (D’Orazio and Green 2021; Raisch and Valkov 2022). Many transcripts increased in abundance when Not1 or Pop2 were depleted, corresponding with the role of CCR4-NOT in initiating the mRNA decay. Many genes also decreased in abundance in response to CCR4-NOT depletion, likely representing secondary consequences downstream of direct CCR4-NOT targets. We observed enrichment of specific classes of CCR4-NOT-affected genes, indicating that one or more regulatory mechanisms coordinate the CCR4-NOT-mediated decay of groups of functionally related transcripts. Future analysis of CCR4-NOT-mediated mRNA metabolism is worth pursuing using new technologies for measuring dynamic changes in poly(A) tail length and RNA metabolism.

Our analysis revealed a functional relationship between codon optimality and Pum activity. mRNAs with low codon optimality are more likely to be susceptible to Pum-mediated repression. We previously observed a similar, albeit weaker, relationship for transcripts degraded by human Pumilio orthologs, PUM1 and PUM2 (Wolfe et al. 2020). In contrast, increased codon optimality correlates with decreased responsiveness to Pum. We observed a similar relationship for the responses of transcripts to CCR4-NOT components Pop2 and Not1. Thus, transcripts with high codon optimality appear to resist the action of decay factors. We speculate that their efficient translation may confer protection (Bae and Coller 2022; Wu and Bazzini 2023). While the molecular mechanism of codon optimality-mediated decay remains to be elucidated, recent evidence from yeast indicates that CCR4-NOT is a central player (Buschauer et al. 2020). In this context, CCR4-NOT senses the empty E-site of poorly elongating ribosomes, leading to subsequent mRNA decay. In contrast, mRNAs with high codon optimality are thought to resist CCR4-NOT due to the relative lack of available empty E-sites during their efficient elongation. Future research is necessary to elucidate the precise mechanism of codon mediated mRNA degradation in metazoans, and our results support that *Drosophila* is a useful model system to do so.

## Acknowledgements

We thank Katherine McKenney, Isioma Enwerem, Robert Connacher, and Elise Dunshee for their insightful comments, suggestions, and technical assistance.

## Contributions

R.J.H., A.C.G. Conceptualization, investigation, resources, formal analysis, writing, review, editing, visualization

A.C.G. Project administration, supervision

C.B., P.L.F. Data analysis, formal analysis, writing, review, editing, visualization, software

P.L.F. supervision

N.D.E., M.K.J., P.J., E.J.W. experiments, data analysis, writing, review

E.J.W. supervision

H.L., H.D.L., C.A.S. investigation, resources, formal analysis, writing, review, data analysis

## SUPPLEMENTAL FIGURES

**Figure S1. Genotype analysis of Pum-myc *Drosophila* DL1 cells.**

**Figure S2. Enrichment sequence motifs in Pum KO + Pum, Pop2, and Not1 RNAi RNA-Seq datasets.**

**Figure S3. Genotype analysis of Pum knockout *Drosophila* DL1 cells, Pum KO1.**

**Figure S4. Identification of differentially expressed genes in response to knockout of Pum in two clonal lines of *Drosophila* DL1 cells.**

**Figure S5. Genotype analysis of V5-Raf in DL1 Pum-myc cells.**

**Figure S6. Gene ontology analysis of differentially expressed transcripts in response to Not1 knockdown relative to negative control RNAi (NTC).**

**Figure S7. Gene ontology analysis of differentially expressed transcripts in response to Pop2 knockdown relative to negative control RNAi (NTC).**

## SUPPLEMENTAL TABLES

**Table S1.** PRE containing transcripts in the *Drosophila* transcriptome

**Table S2.** Click-Seq and differential gene expression data for Pum, Not1, and Pop2 knockdown in DL1 cells.

**Table S3.** Click-Seq and differential gene expression data for Pum knockout and wild type DL1 cells.

**Table S4.** Pum-bound transcripts in *Drosophila* embryos identified by RIP-Seq.

**Table S5.** Intersecting significantly upregulated genes in Pop2, Not1, and Pum knockdown conditions in DL1 cells.

**Table S6.** Drosophila rRNA and 7SL depletion oligonucleotides.

